# Single-Cell Multiomic Analysis of Circadian Rhythmicity in Mouse Liver

**DOI:** 10.1101/2025.04.03.647044

**Authors:** Chun Yip Tong, Changhao Li, Audrey Jacq, Xinyu Y Nie, Chanté R. Guy, Ju Hyun Suh, Raymond K.W. Wong, Christine Merlin, Jerome S Menet, Yuchao Jiang

**Affiliations:** Department of Biology, College of Arts and Sciences, Texas A&M University, College Station, TX, USA; Department of Statistics, College of Arts and Sciences, Texas A&M University, College Station, TX, USA; Center for Biological Clocks Research, Texas A&M University, College Station, TX, USA; Department of Biomedical Engineering, College of Engineering, Texas A&M University, College Station, TX, USA; Center for Statistical Bioinformatics, Texas A&M University, College Station, TX, USA

## Abstract

Circadian rhythms are remarkably widespread across most organisms, regulating hormonal, metabolic, physiological, and behavioral oscillations through molecular clocks that orchestrate the rhythmic expression of thousands of genes. Here, we generate single-nucleus RNA and ATAC multiomics data to simultaneously characterize gene expression and chromatin accessibility of mouse liver cells across the 24-hour day. We interrogate multimodal circadian rhythmicity in both discretized cell types and transient sub-lobule cell states, capturing space-time omics profiles. We delve beyond mean cyclic patterns to characterize stochastic transcriptional bursting and infer spatiotemporal gene regulatory networks that control circadian rhythmicity and liver physiology. Our findings apply to existing single-cell data of mouse and *Drosophila* brains and are validated by time-series single-molecule fluorescence in situ hybridization and vast amounts of orthogonal omics data. Altogether, our study constructs a comprehensive map of the time-series transcriptomic and epigenomic landscapes that elucidate the function and mechanism of the liver peripheral clocks.

## INTRODUCTION

From bacteria to humans, most organisms exhibit inherent 24-hour circadian rhythms ^1^. Best exemplified by the sleep-wake cycle, these rhythms are remarkably widespread, governing hormonal, metabolic, physiological, and behavioral oscillations and enabling biological functions to perform optimally at the most appropriate time of the day ^2–4^. Circadian rhythms are driven by an autonomous and intrinsic timekeeping system known as the circadian clock ^5,6^, which is genetically orchestrated by a set of core clock activator and repressor transcription factors (TFs) interlocked in three transcription-translation feedback loops ^7^. These TFs bind to E-boxes ^8^, ROREs ^9^, or D-boxes ^10^ in the promoters and enhancers of specific target genes to generate cycles of transcription with various phases. Disruption of circadian rhythm has been associated with sleep disorders ^11^, major depression disorder ^12^, cancer ^13^, metabolic syndrome ^14^, Alzheimer’s disease ^15^, and aging ^16^, highlighting the significant role of molecular circadian clocks in physiological processes and disease outcomes.

Recent advances in high-throughput omics technologies provide appealing platforms for identifying circadian genes ^17^ and uncovering the mechanisms that underlie their rhythmic expression ^18^ on the genome-wide scale. In a typical high-throughput experiment, tissues are collected at six to eight different zeitgeber time (ZT) or circadian time (CT) points, followed by omics profiling such as RNA sequencing (RNA-seq) to generate time-series expression profiles. While significant efforts have yielded notable successes – see, for example, existing benchmark studies ^19–21^ and omics databases ^22–24^, bulk-tissue sequencing homogenizes potentially different cell types and cell states, where thousands to millions of cells are pooled with an averaged readout, masking the intricate cellular heterogeneity. Single-cell sequencing, on the other hand, circumvents the averaging artifacts by enabling cellular-level measurements with a much finer resolution.

Several single-cell RNA-seq (scRNA-seq) studies have generated time-series transcriptomic profiles of cells from *Arabidopsis* ^25,26^, mouse suprachiasmatic nucleus (SCN) ^27,28^, mouse clock neurons ^29^, mouse livers ^30^, and *Drosophila* brains ^31^ at different ZT or CT time points. The pioneering studies by Wen et al. ^27^ and Ma et al. ^29^ focused on identifying rhythmic genes within specific cell types or subtypes of the mouse brain. Dopp et al. ^31^ recently investigated cell-type-specific rhythmicity not only across four ZT time points but also under different sleep or wakefulness states of the *Drosophila* brain cells. Expanding beyond discretized cell types and clusters, Droin et al. ^30^ first adopted scRNA-seq to infer circadian rhythmicity in transient cell states of liver hepatocytes. The mouse liver consists of diverse cell types, including hepatocytes, Kupffer cells, fibroblasts, endothelial cells, T cells, and B cells ^32^. Furthermore, the hepatocytes exhibit a distinct spatial trajectory due to the liver’s anatomical compartmentalization into lobular and sub-lobular zonation ^30^, where a gradient of oxygen and nutrient concentrations along the periportal-pericentral axis drives the metabolic and physiological functions of hepatocytes ^33^. This pronounced cell-type and cell-state specificity enables the interrogation of spatiotemporal gene expression and provides deeper insights into gene regulation across both space and time.

Single-cell data is inherently noisy, and treating cells as independent samples inflates the effective sample size and thus the type I error. To address this, existing studies pooled similar cells through averaging as a common approach. However, this common approach largely forfeits the purpose of single-cell sequencing, which characterizes distributions of gene expression, going beyond the mean measurements. The stochastic nature of gene expression, driven by the pervasive phenomenon of transcriptional bursting, has been previously explored in scRNA-seq data ^34,35^. Yet, relating transcriptional bursting to circadian rhythm on the genome-wide scale using time-series single-cell data remains largely underexplored, except for a few studies that adopted single molecule fluorescence in situ hybridization (smFISH) on a limited number of genes ^36,37^.

Beyond transcriptomic studies ^38^, bulk-tissue circadian investigations have successfully examined epigenome ^39,40^, metabolome ^41^, acetylome ^42^, and proteome ^43^, while existing single-cell circadian studies to date ^27,29–31^ have been limited exclusively to the transcriptome. To our best knowledge, no time-series single-cell assay for transposase-accessible chromatin (ATAC) with sequencing data currently exist that temporally profile genome-wide chromatin accessibility, resulting in several missed opportunities. First, the multimodal rhythmicity of chromatin accessibility across gene bodies, regulatory regions such as promoters and enhancers, and TF-binding motifs remains unexplored. Second, genome-wide mapping of cell-type- and cell-state-specific regulatory regions that are linked to rhythmic gene expressions has yet to be systematically conducted. Third, while inferring gene regulatory logic ^44^ involving a target gene, a cis-regulatory element (CRE), and a TF regulator by single-cell omics data has seen rapid methodological development, the application of these approaches to decipher cell-type- and cell-state-specific regulatory networks of circadian gene expression has remained untapped.

In this study, we generate the first single-nucleus RNA and ATAC multiomic data at six different time points across the 24-hour cycle, simultaneously measuring RNA expression of ∼32,000 genes and DNA accessibility of ∼150,000 peaks in ∼25,000 mouse liver cells. We detected circadian rhythmicity in both discretized liver cell types and transient sub-lobule cell states, revealing spatiotemporal RNA and ATAC profiles in a cell-type- and cell-state-specific manner. Moving beyond mean measurements, we characterized gene expression distributions and demonstrated that the fraction of bursting cells closely recapitulate the rhythmicity seen in averaged expression, as traditionally measured. We explored multimodal circadian rhythmicity and examined priming and lagging effects across modalities. For gene regulation, we linked CREs to target genes and further inferred gene regulatory logic involving target gene, TFs, and CREs for circadian control. We validated our findings using time-series smFISH, as well as the vast amounts of existing and orthogonal high-throughput data from chromatin immunoprecipitation followed by sequencing (ChIP-seq), capture Hi-C, and TF knockout experiments. Furthermore, we validated key results in existing scRNA-seq data of mouse and *Drosophila* brains. Altogether, our research constructs a comprehensive map of the transcriptomic and epigenomic landscapes that underpin the function and mechanism of the liver peripheral clocks.

## RESULTS

### Single-nucleus RNA and ATAC multiomic profiling of mouse liver across time points

Using 10x Genomics’ Single Cell Multiome ATAC + Gene Expression solution, we jointly charaterized gene expression and chromatin accessibility of approximately 25,000 mouse liver nuclei, collected at six Zeitgeber time (ZT) points over a 24-hour period under a light-dark experimental design (**Fig. 1A**). Detailed protocols for tissue collection, nuclei isolation, library preparation, and sequencing can be found in the Methods section. Our overall analysis framework is outlined in **Fig. S1**. We began with doublet removal and quality control (QC) using both the RNA and ATAC modalities, resulting in a genome-wide profiling of 16,368 gene expressions and 149,302 peak accessibilities (**Fig. S2**). It is worth noting that while the multiome protocol primarily detects nuclear transcripts, reflecting “nuclear RNA expression,” we use “expression” to lighten the terminology. Using the ATAC fragments, we further derived gene-specific ATAC activity scores ^45^, which quantify the accessibility of gene bodies and promoters, as well as TF-specific motif deviation scores ^46^, which assess the overall accessibility of corresponding TF-binding motifs (**Fig. 1A**).

**Fig. 1.**
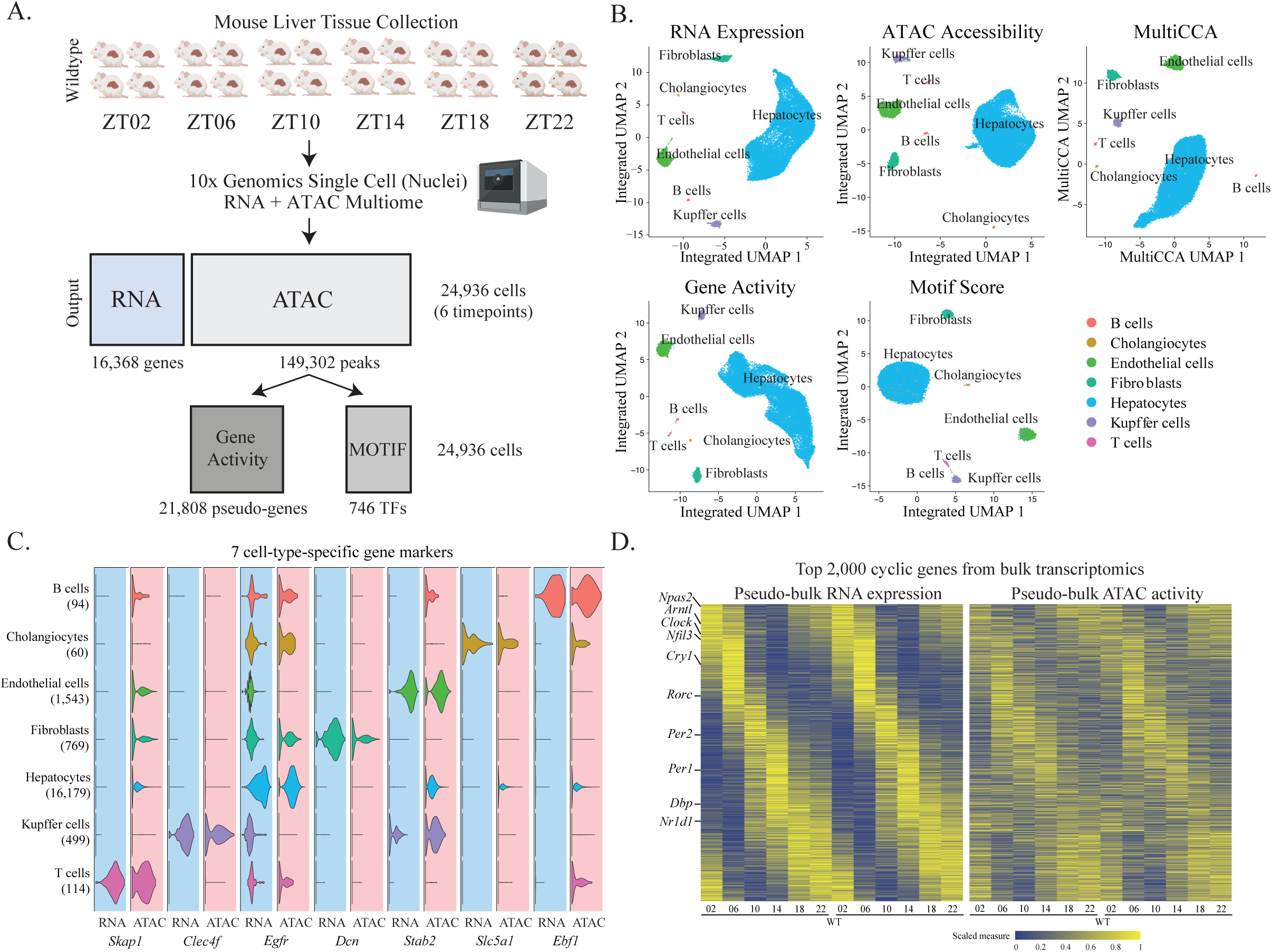
Single-nucleus RNA expression and chromatin accessibility multiomic analysis of mouse liver over a 24-hour cycle. (A) Experimental design. Mouse liver tissues were collected from wild type mice at six time points and processed using the 10x Genomics Chromium Single Cell Multiome platform, enabling simultaneous measurement of RNA expression and chromatin accessibility in the same cells. Liver tissues from four male mice were pooled in equal proportions per time point to minimize inter-mouse variability. (B) Within-modality UMAP dimension reduction of RNA expression, ATAC peak accessibility, ATAC gene activity, motif deviation score, and cross-modality integration using MultiCCA. (C) RNA and ATAC profiles of previously reported cell-type-specific marker genes. (D) Pseudo-bulk RNA and ATAC profiles recapitulate rhythmicity of previously identified circadian genes (top 2000 cyclic genes from Hughes et al. ^49^). All cells were combined to generate pseudo-bulk samples, with average RNA expression and ATAC activity calculated.

Within each of the four modalities (i.e., RNA expression, ATAC accessibility, gene activity, and motif deviation score), we first performed within-modality data normalization, batch-effect correction across time points, and post-correction dimension reduction (**Fig. S3**). This was followed by between-modality data integration via multiple canonical correlation analysis (multiCCA) to borrow information across different modalities ^47^ (**Fig. 1B**). Cell-type labels were transferred from the Liver Cell Atlas (LCA) ^32^ by integrating the RNA domains (**Fig. S4**). This analytical framework effectively identified and distinctly clustered seven major liver cell types: hepatocytes, endothelial cells, fibroblasts, Kupffer cells, cholangiocytes, B cells, and T cells (**Fig. 1B**). Notably, as previously reported by the LCA ^32^, isolation of liver nuclei leads to enrichment of hepatocytes while depleting endothelial cells – a trend we also observed in our data.

As a sanity check, we examined previously reported cell-type-specific marker genes and confirmed that they were enriched within their respective cell types for both the RNA and ATAC modalities (**Fig. 1C**). On the genome-wide scale, cell-type-specific gene expressions from our study closely recapitulated those previously identified by the LCA ^32^, with near-perfect correlation coefficients (**Fig. S5**). Additionally, the ATAC peaks that we detected overlapped with over 90% of the existing bulk-tissue DNase I hypersensitive sites (DHSs) of the mouse liver ^48^. With enhanced cellular resolution, we also detected a signficant number of novel peaks that were validated by histone modifications, serving as candidate CREs (**Fig. S6**).

For circadian rhythm detection, we first sought to reproduce the cyclic patterns previously reported in mouse liver at the bulk-tissue level. To do this, we randomly grouped all cells into two pseudo-bulk samples, analagous to the two replicates in traditional bulk-tissue sequencing. The top 2,000 circadian genes previously identified in mouse liver ^49^ were visualized as heatmaps for both RNA expression and ATAC activity, revealing a clear genome-wide rhythmic pattern in both modalities (**Fig. 1D**). This provides strong empirical support that our generated data can detect genome-wide rhythmicity in chromatin accessibility associated with rhythmic RNA expression. We then trained a deep neural network (DNN) that takes as input gene expression measurements to predict the ZT time point. In the held-out single-cell validation set, the DNN achieved an astounding 99.8% accuracy; in a test set of 1,096 time-series GRO-seq ^50^, Nascent-seq ^51^, and RNA-seq ^51^ samples from 57 public repositories ^38^, the model achieved a strong correlation between the predictions and the ground truths (*r* = 0.84) (**Fig. S7**). This not only provides a versatile tool to make a prediction of time point^52^ but also highlights the high quality of our time-series transcriptomic and epigenomic profiles of mouse liver.

### Cell-type-specific rhythmic RNA expression and ATAC accessibility predominantly in hepatocytes

Using the normalized, batch-corrected, and integrated multi-modal data along with the transferred cell-type labels, we next investigated cell-type-specific rhythmicity across different modalities, focusing on four major cell types – hepatocytes, endothelial cells, fibroblasts, and Kupffer cells (**Fig. 2A**). To ensure an unbiased comparison with an equal testing power across cell types, we downsampled cells from different cell types to retain the same cell count at each time point. Additionally, and more importantly, to strike a balance between type I error control and statistical power and to enhance signal-to-noise ratio in the highly sparse and noisy single-cell data, we pooled similar cells together and constructed metacells at each ZT time point. The number of metacells was determined using gold-standard rhythmic and arrhythmic genes as positive and negative controls ^53^ (**Fig. S8**; see the Method section for more details). Given the time-series omic profiles of the reconstructed metacells, we tested for significant circadian rhythmicity by integrating results from both JTK cycle ^54,55^ and harmonic regression ^56^ via a combination testing framework ^57^, followed by multiple testing correction (**Fig. S9**).

**Fig. 2.**
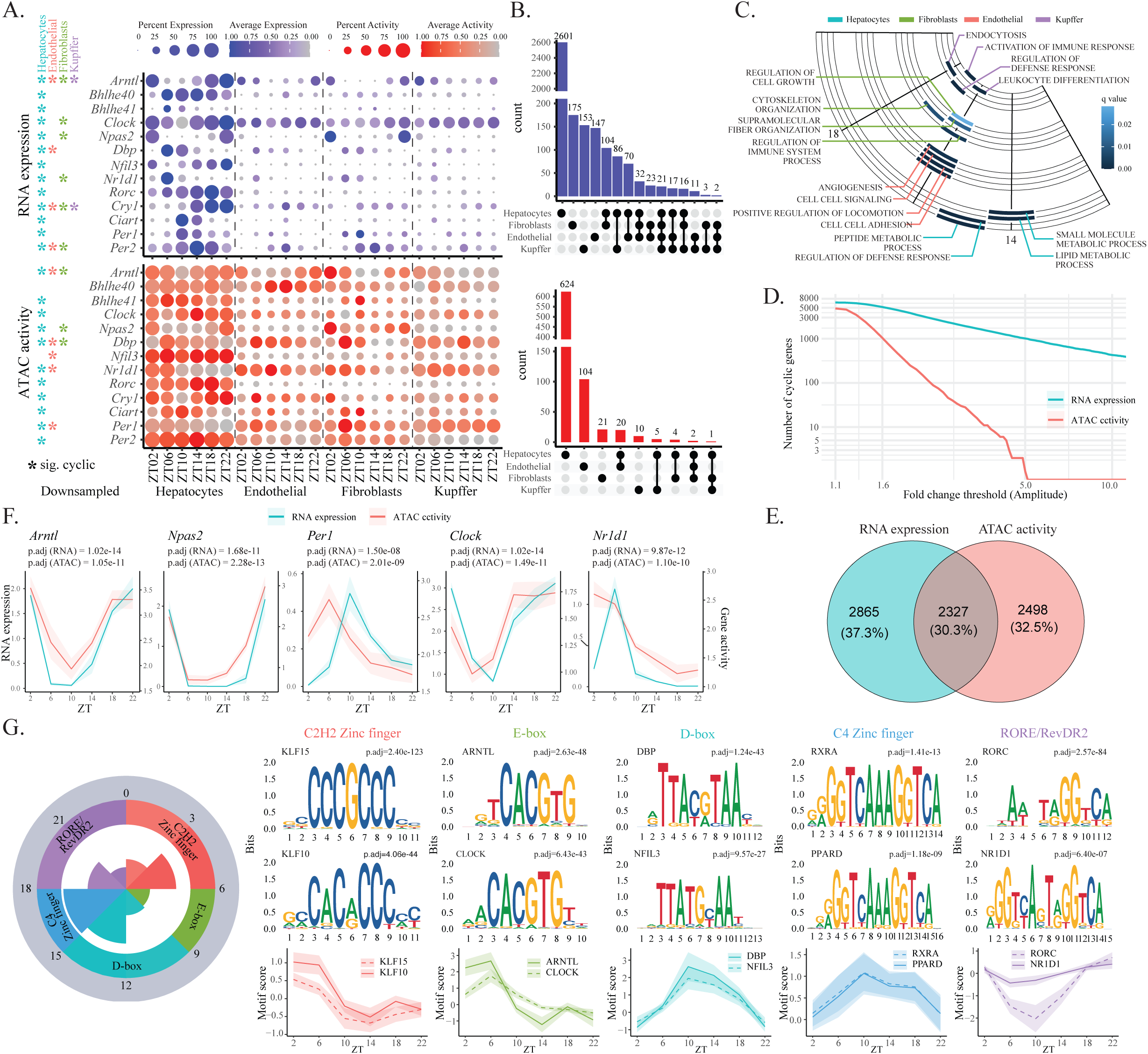
Rhythmic RNA expression and ATAC activity vary across liver cell types and are the most pronounced in hepatocytes. (A) Dot plots show the average RNA expression and ATAC activity (dot color) and the percentage of cells with non-zero RNA expression and ATAC activity (dot size) for core clock and clock reference genes across time points and cell types. Statistically significant rhythmicities are marked with asterisks. To ensure equal testing power, hepatocytes, endothelial cells, fibroblasts, and Kupffer cells were downsampled to the same number of cells. (B) Upset plots depict the gene overlap of cell-type-specific rhythmicity for both RNA expression and ATAC activity, highlighting hepatocytes as the primary contributors to liver rhythmicity. (C) Phase set enrichment analysis identified phase-clustered gene ontologies in different cell types. (D) Cumulative number of rhythmic genes with amplitude fold change above the value on the x-axis for both RNA and ATAC in hepatocytes. (E) Overlap of rhythmic genes between RNA expression (> 1.6 amplitude fold change) and ATAC activity (> 1.1 amplitude fold change) in hepatocytes. (F) Representative clock genes were visualized: *Arntl* and *Npas2* exhibit synchronized rhythmicity between RNA and ATAC modalities, while ATAC activity of *Per1*, *Clock*, and *Nr1d1* shows a significant priming effect relative to their RNA expression. (G) Phase-specific TF-binding motifs significantly enriched in cyclic ATAC peaks. Representative phase-clustered motifs are visualized with position weight matrices and adjusted p-values.

Genome-wide cell-type-specific testing results of circadian rhythmicity are presented in **Table S1**. Examining 13 core clock and clock reference genes (*Arntl*, *Bhlhe40*, *Bhlhe41*, *Clock*, *Npas2*, *Dbp*, *Nfil3*, *Nr1d1*, *Rorc*, *Cry1*, *Ciart*, *Per1*, and *Per2*), we found that nearly all exhibited significant cyclic patterns in hepatocytes for both RNA expression and ATAC activity (**Fig. 2A**). However, in the other cell types (i.e., endothelial cells, fibroblasts, and Kupffer cells), only a subset of the clock genes displayed significant rhythmicity (**Fig. 2A**, **Fig. S10**). For example, *Clock* and *Npas2*, key regulators of the clock through their heterodimeric interactions with *Arntl*, were only significantly rhythmic in hepatocytes and fibroblasts. Consistent with this, a previous study using total RNA-seq confirmed that *Clock* and *Npas2* are not rhythmic in isolated Kupffer cells ^58^. A similar pattern emerged at the genome-wide scale, with hepatocytes harvesting the highest number of uniquely identified rhythmic genes among all cell types (**Fig. 2B**). Notably, hepatocytes also harbor substantially higher numbers of read counts and gene counts than the other liver cell types (**Fig. S11A**), and this observation is consistent with results from the LCA (**Fig. S11B**). While the read counts were normalized by the library size (i.e., the total number of reads) to make them comparable across cell types, this seemingly high difference is also biological – cell size has been shown to correlate with the total number of transcripts and thus the total number of reads per cell ^59^, and hepatocytes are markedly larger than other liver cell types ^60^. Phase set enrichment analysis (PSEA) ^61^ of cell-type-specific rhythmic genes further revealed distinct phase-clustered functional enrichments (**Fig. 2C**, **Fig. S12**). Altogether, our findings reveal significant cell-type-specific rhythmic patterns, aligning with previous scRNA-seq studies of circadian rhythms ^27,29^. In the mouse liver, we demonstrate that hepatocytes – known to play a crucial role in cells’ metabolic functions such as glucose metabolism, lipid metabolism, and detoxification ^62^ – exhibit the most pronounced rhythmic patterns across various modalities of the peripheral clock.

Moving beyond RNA expression to compare across modalities, we first plotted the cumulative number of rhythmic genes at varying amplitude fold change thresholds for both RNA and ATAC (**Fig. 2D**). Applying a threshold of RNA expression fold change > 1.6 and ATAC activity fold change > 1.1, we identified 2,865 significantly rhythmic genes in RNA expression and 2,498 in ATAC activity, with 2,327 genes exhibiting rhythmicity in both; these results were obtained from the full set of hepatocytes, considering 15,159 genes after RNA and ATAC gene QC (**Fig. 2E**). For genes that are rhythmic for both modalities, their phases of RNA expression and ATAC activity are highly correlated (*r* = 0.83; **Fig. S13A**). We further tested for significant phase shifts between the two modalities using CircaCompare ^63^. Clock genes such as *Arntl* and *Npas2* exhibit synchronized rhythmicity, while the ATAC activity of *Per1*, *Clock*, and *Nr1d1* harbors an earlier phase than RNA expression, suggesting a temporal relationship between chromatin state and transcription (**Fig. 2F**). On a genome-wide scale, genes with significantly advanced ATAC phases (i.e., those where ATAC rhythmicity primes RNA rhythmicity) are longer and exhibit higher relative amplitude (rAMP) (**Fig. S13B**), potentially due to their greater need for chromatin priming. Notably, for genes with advanced RNA phases, while their precise regulatory mechanism requires further investigation beyond the scope of this paper, a recent study ^64^ suggested that chromatin remodeling can occurs four hours after transcription – the translational process facilitates chromatin remodeling ^64^ and meanwhile transcribed RNAs form R-loop structures with CREs to promote recruitment of epigenetic modifiers ^65^.

Next, we analyzed 746 TFs with annotated binding motifs from the JASPAR database ^66^ and found that 20% of these TFs exhibited significant rhythmicity in gene expression, ATAC activity, and motif scores (**Fig. S14A**), with genome-wide results shown in **Table S2**. Notably, differences between TF gene expressions and TF motif scores likely result from their distinct measures of transcription and DNA binding. For TF-binding motifs, deviation scores of E-box motifs (ARNTL, CLOCK, BHLHE40, and BHLHE41), D-box motifs (DBP and NFIL3), and RORE motifs (NR1D1 and RORC) display distinct phases, demonstrating their collective function to regulate rhythmic gene expression around the clock (**Fig. S14B**). Importantly, TF-specific rhythms were also more pronounced in hepatocytes than in the other cell types – some exhibited unique circadian patterns, while others showed heightened relative amplitudes (**Fig. S14B**) – aligning with our previous observations based on the gene-level testing results.

In addition to aggregating ATAC fragments into either gene activity or motif deviation scores, we identified 15,665 ATAC peaks (10.5% of ∼150K peaks) with significant rhythmic accessibility. This contrasts with a previous bulk-tissue study, which identified 8% of ∼65K DHSs as cyclic ^67^, highlighting the enhanced resolution and increased testing power. We further carried out a phase-specific motif enrichment analysis using the significantly cyclic peaks (**Fig. 2G**). We discovered enrichment of E-box motifs from ZT06 to ZT09, D-box motifs from ZT09 to ZT15, as well as RORE and RevDR2 motifs from ZT18 to ZT24, consistent with a previous report ^50^. Interestingly, we also found that the binding motifs for KLF15 and KLF10 were highly enriched between ZT00 and ZT06. The Krüppel-like family of transcription factors (KLFs) are known to play essential roles in liver physiology and metabolism ^68^; previous research has shown that *Klf15* and *Klf10* expression is strongly induced in the liver during fasting, aligning with the observed enriched timeframe ^69^. Additionally, the binding motifs of Retinoid X Receptor Alpha (RXRA) and Peroxisome Proliferator-Activated Receptor Delta (PPARD) were highly enriched between ZT15 and ZT18; both genes play critical roles in metabolic processes linked to the feeding cycle, functioning as a heterodimer to regulate lipid metabolic pathways ^70^.

### Beyond mean measurements: rhythmic nuclear gene expression driven by rhythmic bursting fraction

Single-cell sequencing circumvents the averaging artifacts associated with traditional bulk-tissue sequencing, thus enabling characterization of not only mean measurements but also distributions. To date, several scRNA-seq studies ^34,35^ have demonstrated that cellular gene expression is highly stochastic, resulting from the pervasive phenomenon of transcriptional bursting, where cells switch between “ON” and “OFF” transcriptional states. While pioneering studies have adopted time-series smFISH to study circadian rhythmicity in relation to promoter-enhancer looping and transcriptional bursting ^30,37,71^, they focused on a limited set of clock genes (e.g., *Arntl*, *Nr1d1*, and *Cry1*, as shown in **Fig. S15**), leaving behind a missed opportunity for comprehensive genome-wide profiling.

To systematically investigate the relationship between transcriptional bursting and circadian rhythm, we tested two different underlying models that can both result in cyclic gene expressions: rhythms in average expression across the majority of cells (Model 1) and rhythms in the fraction of bursting cells (Model 2) (**Fig. 3A**). For model 1, we calculated the average expression across all cells comprising the metacell, while for model 2, we computed the fraction of cells with non-zero expression. Both the average expression and the bursting-cell fraction measured at the metacell level at different time points were used as input to detect circadian rhythmicity. Note that these two measures differ, as single-cell gene expression measurements from unique molecular identifier (UMI)-based protocols have been shown to follow a Poisson distribution ^72^. Without loss of generality, we focused on hepatocytes and later extended the testing framework to other cell types, tissues, and species. Interestingly, in hepatocytes, all core clock and clock reference genes – including *Arntl*, *Per1*, *Per2*, *Npas2*, *Dbp*, *Cry1*, *Clock*, *Nr1d1*, and *Rorc* – exhibited significant rhythmicity for both mean RNA expression and the proportion of cells with non-zero expression (i.e., the fraction of bursting cells), with nearly identical cosinor curves (**Fig. 3B**). Our results strongly suggest that the mean of binarized expression (i.e., the fraction of cells with non-zero expression) closely mirrors the mean of the absolute integer-valued expression in the circadian context. Biologically, this indicates that the fraction of cells actively transcribing RNAs varies between the peak and trough, regardless of the burst magnitude or the quantity of RNAs produced. This observation, combined with the widespread monoallelic expression observed at the single-cell level ^73,74^ due to transcriptional bursting, overwhelmingly supports model 2 over model 1.

**Fig. 3.**
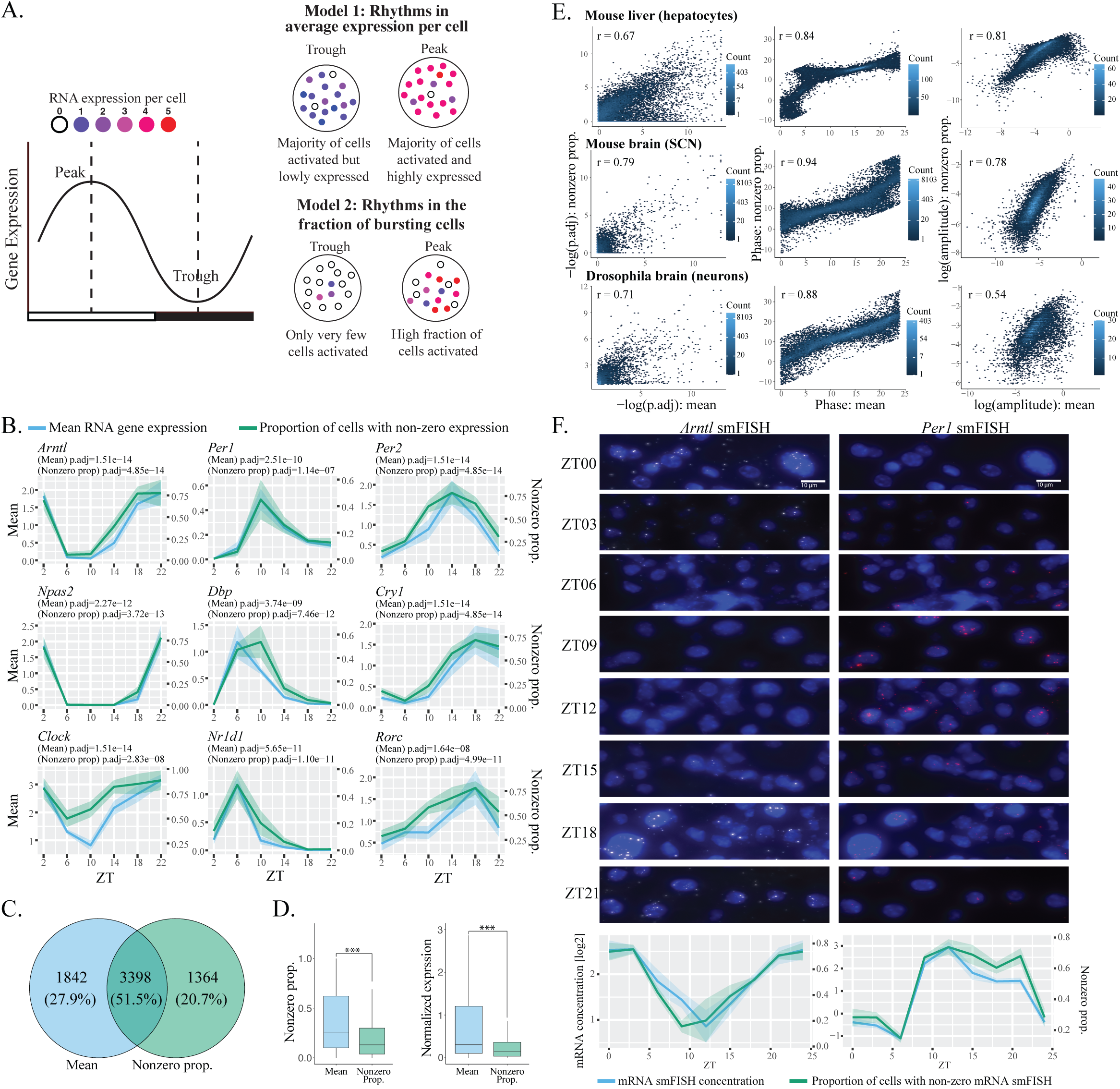
Moving beyond mean measurements: rhythmic nuclear gene expression driven by rhythmic bursting fraction. (A) Two models of circadian rhythmicity: one driven by the average expression across cells and the other by the fraction of bursting cells with non-zero transcripts. (B) Core clock and clock reference genes exhibit nearly identical cyclic patterns of both mean expression and bursting fraction. (C) Venn diagram depicting the gene overlap between rhythmicity detected by mean expression and bursting fraction. (D) Boxplot of RNA expression for cyclic genes identified by mean expression and bursting fraction. Cyclic genes identified by mean expression have significantly higher fractions of cells in the “ON” bursting state and significantly higher absolute expression levels. (E) Correlations of adjusted p-values (left), amplitudes (middle), and phases (right) between rhythmicity detected by mean expression and bursting fraction. Results are reproduced using time-series scRNA-seq data from mouse SCN ^27^ and *Drosophila* neuron ^29^. (F) Orthogonal and experimental validation by smFISH of *Arntl* and *Per1* in mouse liver at different time points ^30^. Mean RNA concentration and proportion of nuclei with non-zero transcripts show near-identical cyclic patterns.

Extending the investigation to the genome-wide scale, 51.5% of rhythmic genes showed significant rhythmicity in both mean expression and bursting-cell fraction (**Fig. 3C**). Of the 1,842 genes with only rhythmic mean expression and the 1,364 genes with only rhythmic bursting-cell fraction, the former group shows significantly less “burstiness” and higher expression levels (**Fig. 3D**). The bursting kinetics of constitutively expressed genes (e.g., housekeeping genes) become statistically intractable due to identifiability issues, as previously reported ^34,75^. Therefore, due to the lack of burstiness, accurately inferring their bursting parameters when fitting the Poisson-Beta hierarchical model is neither possible nor necessary. When comparing rhythmic genes identified based on the mean expression, the proportion of cells with non-zero expression (a surrogate for burst frequency), and the mean of non-zero expression (a surrogate for burst size), we found significant overlap between the first two groups of genes, whereas far fewer genes were detected solely by non-zero mean expression (**Fig. S16A**). Additionally, Gini index and coefficient of variation – both measuring variability – were almost perfectly correlated with mean expression and the fraction of bursting cells (**Fig. S16B**), indicating that biological variation also exhibits significant rhythmic patterns.

For reproducibility, both gene-specific and genome-wide results apply to existing time-series scRNA-seq data of mouse SCN ^27^ (**Fig. S17**) and *Drosophila* neuron ^29^ (**Fig. S18**). Across all datasets, adjusted p-values, phases, and amplitudes showed strong correlations between testing of cyclic mean expression and cyclic bursting-cell fraction (**Fig. 3E**). For validation, we utilized previously published smFISH data with both mean mRNA concentration and proportion of cells with detectable transcripts in the nuclei for *Arntl* and *Per1* ^30^ (see Method for detail). Consistent with our single-nucleus sequencing results, the rhythmic pattern of the bursting-cell fraction closely mirrors that of the mean measurement – the nuclear RNA concentration calculated across cell nuclei on a log scale for variance stabilization (**Fig. 3F**). Collectively, our results, overwhelmingly supported by empirical evidence from extensive orthogonal and independent omics resources, suggest that circadian rhythmicity is predominantly driven by burst frequency rather than burst size. We further relate this discovery and conclusion to other biological contexts in the Discussion section.

### Spatiotemporal gene expression and CRE accessibility in hepatocytes across the liver lobules

So far, our analyses have focused on discretized cell types, while the mammalian liver also comprises transient cell states due to its hexagon-shaped lobules, with central vein (CV) at the center and portal triad at the corners (**Fig. 4A**). The portal triad consists of a portal node (PN), portal artery, and bile duct. Blood flows from the PN towards the CV through sinusoids, creating a directional gradient of oxygen, nutrients, and hormones ^76,77^. To infer spatial zonation ^30,33^ of our sequenced cells along the lobule axis, we performed trajectory reconstruction using Slingshot ^78^ and inferred pseudotime, a fractional measure capturing cells’ spatial positioning along the PN-to-CV axis (**Fig. 4B**). As expected, previously reported ^33^ gene markers *Cyp2f2* and *Cyp2e1* were predominantly expressed and accessible in periportal and pericentral hepatocytes near the PN and CV, respectively (**Fig. 4C**).

**Fig. 4.**
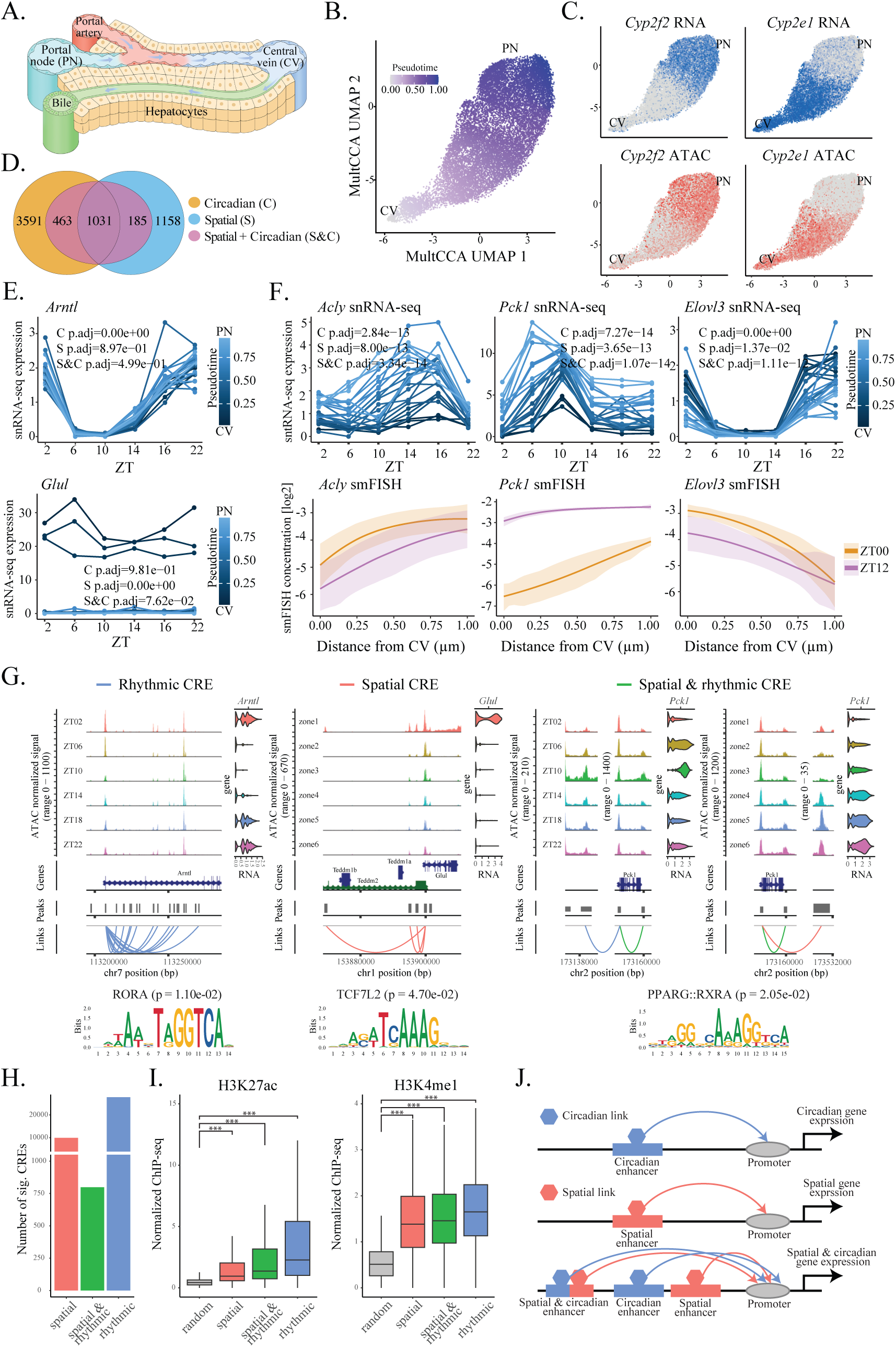
Spatiotemporal gene expression and regulation across lobular zonation and time points. (A) Schematic of the liver lobule, where concentric layers of hepatocytes are positioned along PN-to-CV axis. Blood flow and secreted morphogens give rise to a spatially graded microenvironment, resulting in different functions assigned to different zonation. (B) UMAP dimension reduction of hepatocytes colored by inferred pseudotime. Spatial distribution of RNA expression and ATAC activity for two marker genes, *Cyp2f2* and *Cyp2e1*, enriched at the PN and CV, respectively. (D) Venn diagram of circadian, spatial, and spatial and circadian genes detected. (E) Visualization of a circadian gene *Arntl* and a spatial gene *Glul*. (F) Visualization of spatial and circadian genes, validated by smFISH. (G) Representative CREs regulating circadian and/or spatial expression of target genes. Cells are grouped by either time points or spatial zonation. Enriched motifs are identified from each target genes’ significantly linked CREs and are consistent with previous reports. (H) Number of CREs that regulate circadian rhythms, spatial expression, or both simultaneously. (I) ChIP-seq of H3K27ac and H3K4me1 ^82^, active enhancer markers, for rhythmic, spatial, and combined spatial and rhythmic CREs, compared to random regions from the mouse genome. (J) Illustration of CREs significantly linked to target genes regulating circadian rhythmicity, spatial variation, or both.

Given the time points and the inferred spatial zonation of the cells, we devised a statistical framework to test whether each gene’s RNA expression and ATAC activity exhibit spatial, circadian, or combined spatial and circadian patterns (**Fig. 4D**; see Method for details and **Fig. S19** for testing results of representative genes). Our results revealed that the core clock gene *Arntl* was circadian but not spatial, ensuring a robust circadian control across the hepatic zonation ^30^ (**Fig. 4E**). The other clock reference genes all exhibited significant cyclic patterns, although some were also spatially variable, especially when jointly tested across both time and location (**Fig. S20**). In contrast, *Glul*, which was previously shown to be spatially enriched at the CV ^76^, was spatially variable but not circadian (**Fig. 4E**). At the genome-wide level, we identified transient genes with spatially variable RNA expression and/or ATAC activity enriched near the PN and CV, followed by KEGG pathway enrichment analysis (**Fig. S21**). For spatiotemporal expression, we identified 1,679 genes across the genome that were both spatially variable and rhythmic over time. Importantly, the joint testing framework across both spatial location and time point identified spatial and circadian genes that were otherwise masked when testing by location or time point alone, highlighting the utility of the spatiotemporal approach (**Fig. S19**). **Fig. 4F** highlights three representative spatial and circadian genes – *Acly*, *Pck1*, and *Elovl3* – that exhibit significant space-time variation. Their spatiotemporal patterns were further validated by time-series smFISH of mouse liver ^30^. Notably, beyond RNA expression, ATAC activity of genes also demonstrated similar spatiotemporal patterns (**Fig. S22**).

The liver’s peripheral clock is governed by intricate gene regulatory networks. While previous studies have demonstrated the circadian and transient expression of genes along the liver lobule ^30^, the underlying gene regulatory mechanisms remain largely unexplored. Utilizing the paired ATAC modality, we linked open-chromatin regions (i.e., CREs) to target gene expressions by testing for significant association between chromatin accessibility and gene expression (see the Method for details). Importantly, this association was tested across both spatial and temporal dimensions, allowing us to identify spatial, rhythmic, and spatial & rhythmic CREs. Three representative CRE-gene pairs were visualized in **Fig. 4G**. Specifically, *Arntl*, a core clock gene, exhibited significant temporal correlations with rhythmic CREs. Motif enrichment analysis using the linked CREs returned significant enrichment of the RORA motif, consistent with the role of the ROR family as key regulators of *Arntl* ^79^ (**Fig. 4G**). Additional rhythmic CREs were also identified for other clock genes, including *Clock*, *Cry1*, *Dbp*, *Nr1d1*, and *Rorc* (**Fig. S23**). In contrast, *Glul*, the previously identified spatially variable gene, was significantly associated with spatial CREs enriched for the TCF7L2 binding motifs ^80^ (**Fig. 4G**). Interestingly, *Pck1*, a circadian and transient gene, displayed spatial, temporal, and spatiotemporal correlations with CREs enriched for the binding motif of PPARG and RXRA; these TFs form a heterodimer to regulate the expression of *Pck1*, which is essential for the maintenance of lipid metabolism ^81^ (**Fig. 4G**).

Genome-wide, we identified over 27,500 rhythmic CREs, 9,500 spatial CREs, and 750 spatial & rhythmic CREs (**Fig. 4H**). Detailed linkage results are presented in **Table S3**, with top significantly enriched motifs from the linked CREs displayed in **Fig. S24**. For validation, we utilized existing ChIP-seq data for the enhancer histone markers H3K27ac and H3K4me1 ^82^ enriched at active and primed enhancers; both markers showed significant enrichment at spatial, rhythmic, and spatial & rhythmic CREs, compared to random genomic regions (**Fig. 4I**). By leveraging paired RNA and ATAC modalities characterized at the single-cell level, we were able to identify different types of CREs linked to the same target genes across time points and lobular zonation, with corresponding motif enrichments for clock and liver-specific metabolic TFs (**Fig. 4J**). These genome-wide results not only provide strong empirical support for the classic model of an integrative regulatory mechanism involving multiple CREs with TF-binding motifs ^83,84^ but also reveal spatiotemporally resolved gene regulatory logic across both time and space.

### Three-way relationships between CREs, TFs, and target genes for circadian regulation

Understanding how TFs interact with CREs to regulate target gene expression is essential for elucidating the molecular mechanisms for circadian control. Traditional approaches adopted ChIP-seq to detect TF binding at potential CREs ^85^. Target genes are, however, not directly inferred and are typically assigned based on their proximity to the CREs, often limiting the focus of CREs primarily to the genes’ promoter regions. While the previous section linked more distant CREs to target genes, CRE-gene linkage analysis does not provide direct information on TF binding either – motif enrichment analysis using these linked CREs is a two-step approach that further suffers from reduced resolution of TF binding. In this regard, the single-cell multiomic design enables time-series characterizations of CRE accessibility, TF binding, and target gene expression, all measured simultaneously in the same cells, offering unprecedented opportunities to directly infer regulatory relationships among these factors.

Leveraging the recently developed TRIPOD ^44^ framework – a computational approach for detecting three-way regulatory relationships among TFs, CREs, and target genes using single-cell multiomics data – we next sought to detect regulatory “trios,” where a TF binds to a CRE to regulate target gene expression (**Fig. 5A**). Specifically, TRIPOD takes as input the motif score of a TF (replacing the default TF expression to more accurately capture binding activity and regulatory potential), the accessibility of a candidate CRE, and the expression of a gene to infer the CRE-TF-gene relationship, with genome-wide results shown in **Table S4**. **Fig. 5B** outlines three significant trios that are in line with the transcriptional and translational feedback loops (E-box, D-box, and RORE) of the circadian clock ^7^ and the KEGG mouse circadian rhythm pathway. Specifically, ARNTL and CLOCK form a heterodimer that binds to the E-box in the CREs of *Per1*, activating its transcription ^86^. RORA acts as a transcriptional activator by binding to the RORE motifs to regulate the expression of *Npas2* ^87^. Furthermore, DBP, a PAR bZip TF, activates transcription of *Per* genes through its binding to the D-box in their promoter regions, consistent with a previous report ^88^.

**Fig. 5.**
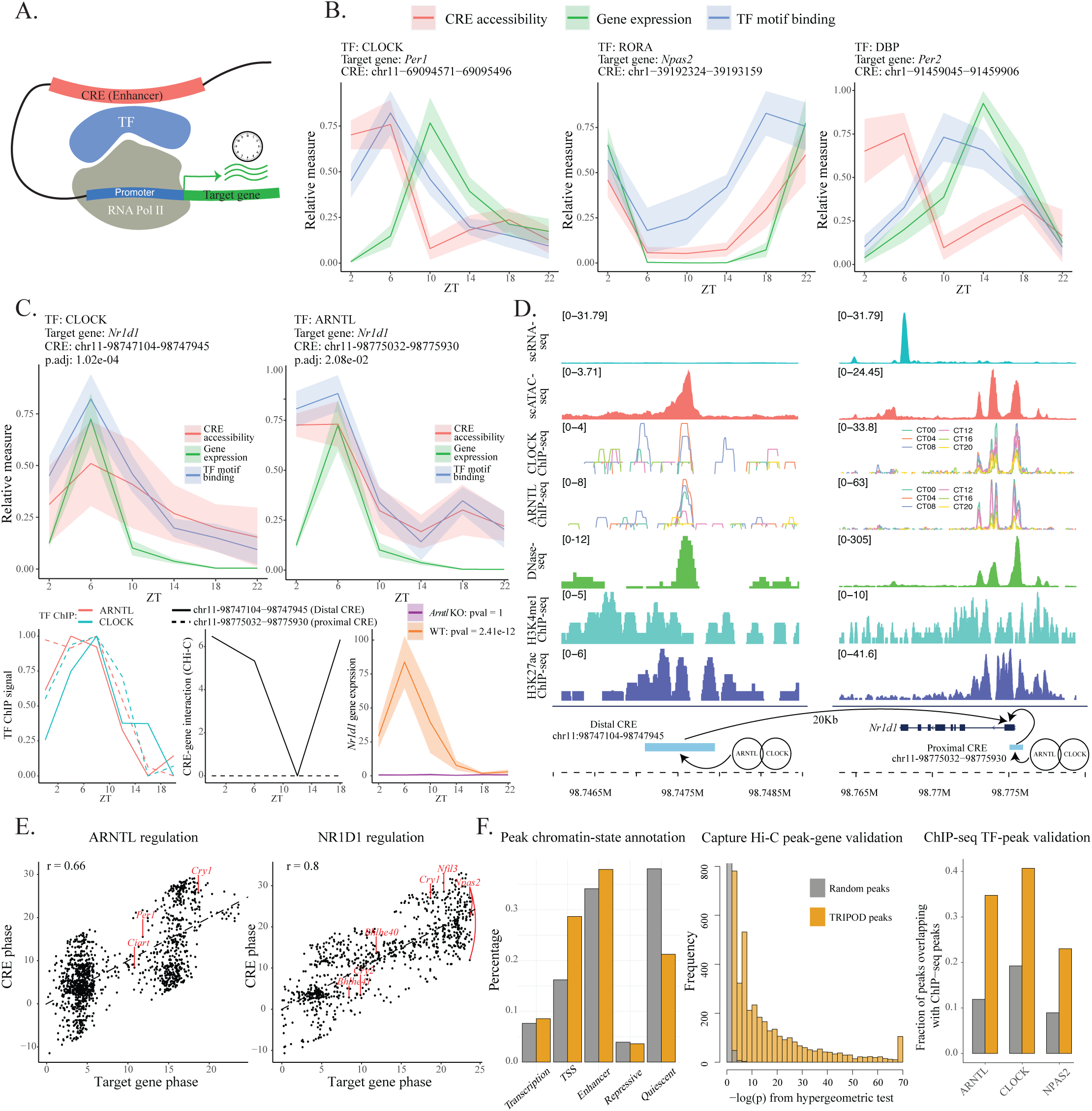
Transcriptional control of rhythmic gene expression in hepatocytes. (A) Schematic of a CRE-TF-gene trio: a TF binds at a CRE to regulate target gene expression. (B) Significant trios consistent with prior knowledge, including the transcription-translation feedback loops for circadian control and the mouse circadian KEGG pathway. (C) ARNTL and CLOCK bind to CREs to regulate *Nr1d1* gene expression ^89^. Validation includes time-series ChIP-seq data confirming ARNTL and CLOCK binding at the CREs ^90^, as well as time-series capture Hi-C data confirming CRE-*Nr1d1* interaction ^91^. *Nr1d1* expression and rhythmicity are depleted upon *Arntl* knockout. (C) Zoomed-in coverage plots of single-nucleus RNA-seq and ATAC-seq, TF ChIP-seq (ARNTL and CLOCK), DNase-seq, and histone ChIP-seq at the predicted proximal and distal CREs and the target gene. (E) Trios involving ARNTL (left) and NR1D1 (right) as TFs. While phases of the target genes and their CREs are highly correlated, they can be out-of-phase with their regulating TFs. This aligns with findings that enhancer RNAs play an essential role in controlling target gene rhythmicity, which is anti- or out-of-phase with TF binding rhythmicity ^50^. (F) Chromatin-state annotation of linked CREs, capture Hi-C validation of CRE-gene interactions ^91^, and ChIP-seq validation of CRE-TF linkage ^90^. Compared to random genomic regions, identified CREs are enriched at TSSs and enhancers, exhibit statistically significant interactions with target genes, and show a markedly higher frequency of TF binding.

We further zoomed into two trios consisting of ARNTL and CLOCK as the TFs, *Nr1d1* as the target gene, and two CREs located at the promoter and ∼20Kb downstream of *Nr1d1* (**Fig. 5C**). While ARNTL and CLOCK have been shown to regulate *Nr1d1* ^89^, we pinpointed a proximal and a distal CRE, in which the TFs bind at the E-box element, respectively. ChIP-seq of ARNTL and CLOCK from mouse liver ^90^ confirmed their binding at both CREs, exhibiting synchronous circadian patterns aligned with the identified trio (**Fig. 5C**). Capture Hi-C ^91^ further confirmed a significantly rhythmic interaction between the distal CRE and the target gene *Nr1d1*, while resolution was limited for the proximal CRE due to its location within the gene’s promoter region (**Fig. 5C**). Moreover, *Nr1d1*’s expression and rhythmicity were both depleted when ARNTL was knocked out ^92^ (**Fig. 5C**). Zoomed-in coverage plots of the *Nr1d1* and CRE regions are shown in **Fig. 5D**, revealing that the predicted CREs were highly accessible, bound by the identified TFs, and enriched with active histone marks. Additionally, our results suggest that *Arntl* itself is regulated by multiple TFs, including MEF2D ^93^, PPARD ^70^, NR5A2 ^91^, and NFYB ^94^, consistent with previous reports (**Fig. S25**). These findings underscore the complex and multifaceted regulatory network governing clock gene expression, highlighting their critical role in circadian rhythm regulation.

Further examination of the identified CRE-TF-gene trios reveals a strong correlation between the phases of CRE accessibility and the phases of the target gene expression, although both can be out of phase with the TF regulator. <u>T</u>wo sets of trios with ARNTL and NR1D1 as the TF regulators are provided in Fig. 5E. We show that the binding of the TF regulators is asynchronous with the expression of the target genes they regulate, consistent with previous reports ^50,95^. By incorporating the additional ATAC modality, we provide evidence that the target gene expression, in addition to being regulated by TFs, is also governed by linked CREs with synchronized accessibilities (**Fig. 5E**). This suggests a finely tuned regulatory system, where both TFs and chromatin states coordinate to ensure precise circadian gene expression control.

Gene regulatory networks for the core clock and clock reference genes were further constructed, including the E-box binding TFs (ARNTL, CLO<u>C</u>K, and NPAS2), the RORE binding TFs (RORA, RORC, and NR1D1), the D-box binding TF (DBP), and TFs without annotated binding motifs (PER1, PER2, and CRY1) (**Fig. S26**). The reconstructed regulatory network is highly consistent with the KEGG circadian rhythm pathway, and we further validated it using chromatin-state annotation, promoter capture Hi-C, and TF ChIP-seq (**Fig. 5F**). Specifically, the predicted CREs were enriched at TSSs and enhancers but depleted in quiescent regions compared to random peaks. Hypergeometric tests demonstrated that the identified CREs were significantly enriched for peak-gene interactions. Furthermore, TFs that are predicted to bind at CREs showed significantly higher ChIP-seq signals for ARNTL, CLOCK, and NPAS2 compared to random peaks.

## DISCUSSION

Recent advances in next-generation sequencing technologies offer appealing platforms to identify circadian genes and elucidate the mechanisms that regulate their rhythmic expression ^17,18^. Traditional bulk-tissue sequencing pools thousands to millions of cells with an averaged readout, masking the intricate cellular diversity by blending various cell types and states. To date, several scRNA-seq studies have measured cellular gene expression at different time points across the 24-hour cycle ^25–31^. However, to the best of our knowledge, this study is the first to generate a single-cell multiomics dataset that simultaneously measures gene expression and chromatin accessibility to investigate circadian rhythmicity. This new technology brings distinct advantages, poised to challenge the existing status quo.

Specifically, this time-series single-cell multiomic design allows us to systematically characterize and disentangle the different levels of heterogeneity: between discretized cell types and transient cell states, across time points and spatial locations, and among different genomic modalities simultaneously measured within the same cells. Mouse liver comprises diverse cell types such as hepatocytes, Kupffer cells, fibroblasts, endothelial cells, T cells, and B cells ^32^. Moreover, the hepatocytes exhibit pronounced subtype specificity due to the liver’s anatomical compartmentalization into lobules ^76^. We identified cell-type-specific rhythmic patterns across different omic modalities and demonstrated that hepatocytes displayed the most pronounced and robust cyclic patterns. Additionally, we examined omics profiles across spatial zonation and revealed not only circadian but also spatial and spatiotemporal gene expressions and regulatory relationships. Notably, while we computationally reconstructed the PN-to-CV trajectory using well-established methods, we did not properly account for the inference uncertainty. Looking ahead, applying recently developed imaging- and sequencing-based spatial transcriptomics technologies – particularly those with subcellular resolution, such as 10x Genomics’ Xenium and Visium HD platforms – will allow direct and unbiased measures of the spatial location.

We demonstrate on the genome-wide scale that the fractions of bursting cells (i.e., cells with non-zero expressions in the snRNA-seq) closely recapitulates the rhythmic patterns of the averaged expressions, as traditionally measured in bulk RNA-seq. This regulation by burst frequency, instead of burst size, has previously been shown to take effect in core clock genes ^36,71^, as well as in several other contexts: across different genes ^96^, during the cell cycle ^97^, under different growth condition ^98^, through regulation by TFs ^99^, and via looped contacts between enhancers and promoters ^71,100^. Importantly, we show that the bursting fractions are strongly correlated with the Gini index and the coefficient of variation, suggesting that “noise” or variation resulting from stochastic transcription also exhibits robust rhythmicity. It is worth noting that while a two-state Poisson-Beta model ^34,75^ has been proposed to infer the bursting kinetics, in this study, we resorted to the fraction of bursting cells as a surrogate measure of burst frequency and the non-zero mean expression as a surrogate measure of burst size. This is primarily due to the highly sparse and noisy data from 10x Genomics’ 3’ end tagging, which led to poor model fitting with identifiability issues ^34,101^. The inherent heterogeneity within the same cell type (e.g., across different spatial zonation and time points) further exacerbated the problem, necessitating a large number of cells for accurate parameter estimation. With recent developments of full-transcript scRNA-seq ^102,103^, we anticipate more accurate estimates of both burst frequency and burst size, potentially at isoform and allele resolution, which will help further disentangle the various components of circadian regulation by transcriptional bursting.

To infer the transcriptional regulation of circadian gene expression, we conducted a genome-wide scan, focusing on one CRE-TF-gene trio at a time. It is important to note that this CRE-TF-gene conditional association does not imply causation, nor does it account for other TF regulators or mediators that may also be involved in this complex regulatory network. With increased data resolution and cell numbers, it would be meaningful to explore beyond the marginal and trio relationships to incorporate higher-order models possibly with variable selection that can more realistically capture the complex regulatory relationships between multiple CREs, modules consisting of multiple TFs, and target gene expression. The abundance of data from past and ongoing bulk sequencing studies, coupled with the maturation of single-cell sequencing protocols, can also be further exploited via a joint deconvolution framework ^104^ that utilizes cellular level rhythmic profiles to uncover cell-type and cell-state contributions in circadian biology.

Last but not least, an interesting observation we want to highlight is that our ATAC fragments showed significant TSS enrichment from ZT02 to ZT06 (**Fig. S27**). While TSS enrichment is typically a QC metric to filter out cells that have failed library preparations and sequencing ^105^, validation using published ChIP-seq data of RNA polymerase II (Pol II), TATA-binding protein (Tbp, responsible for Pol II loading and initiation ^106^), Negative Elongation Factor A (Nelf-A, responsible for stablizing Pol II ^106^), and H3K4me3 histone modification confirmed heightened TSS enrichment at ZT02 (**Fig. S27**). These findings provide deeper insights into the complex role of RNA polymerase II pausing in regulating the temporal dynamics of gene transcription in the mouse liver ^106,107^.

## ACKNOWLEDGEMENTS

This work was supported by the National Institute of Health (NIH) Grants R01 DK128133 (to J.M.), R01 GM145737 (to J.M.), and R35 GM138342 (to Y.J.). The authors thank Drs. Dingbang Ma and Michael Roshbash for providing support on the *Drosophila* neuron data, Drs. Shao’ang Wen and Jun Yan for providing support on the mouse SCN data, Drs. Clémence Hurni and Felix Naef for providing support on the mouse liver smFISH data, Chaohua Wu for helping with ImageJ, and High Performance Research Computing at Texas A&M University for providing computational resources and support.

## AUTHOR CONTRIBUTIONS

J.M. and Y.J. initiated and envisioned the study. A.J., X.N., C.R.G., and J.M. performed the mouse experiments. All authors performed data analysis and results interpretation. C.Y.T. and Y.J. wrote the manuscript, which was reviewed and approved by all authors.

## DATA AVAILABILITY

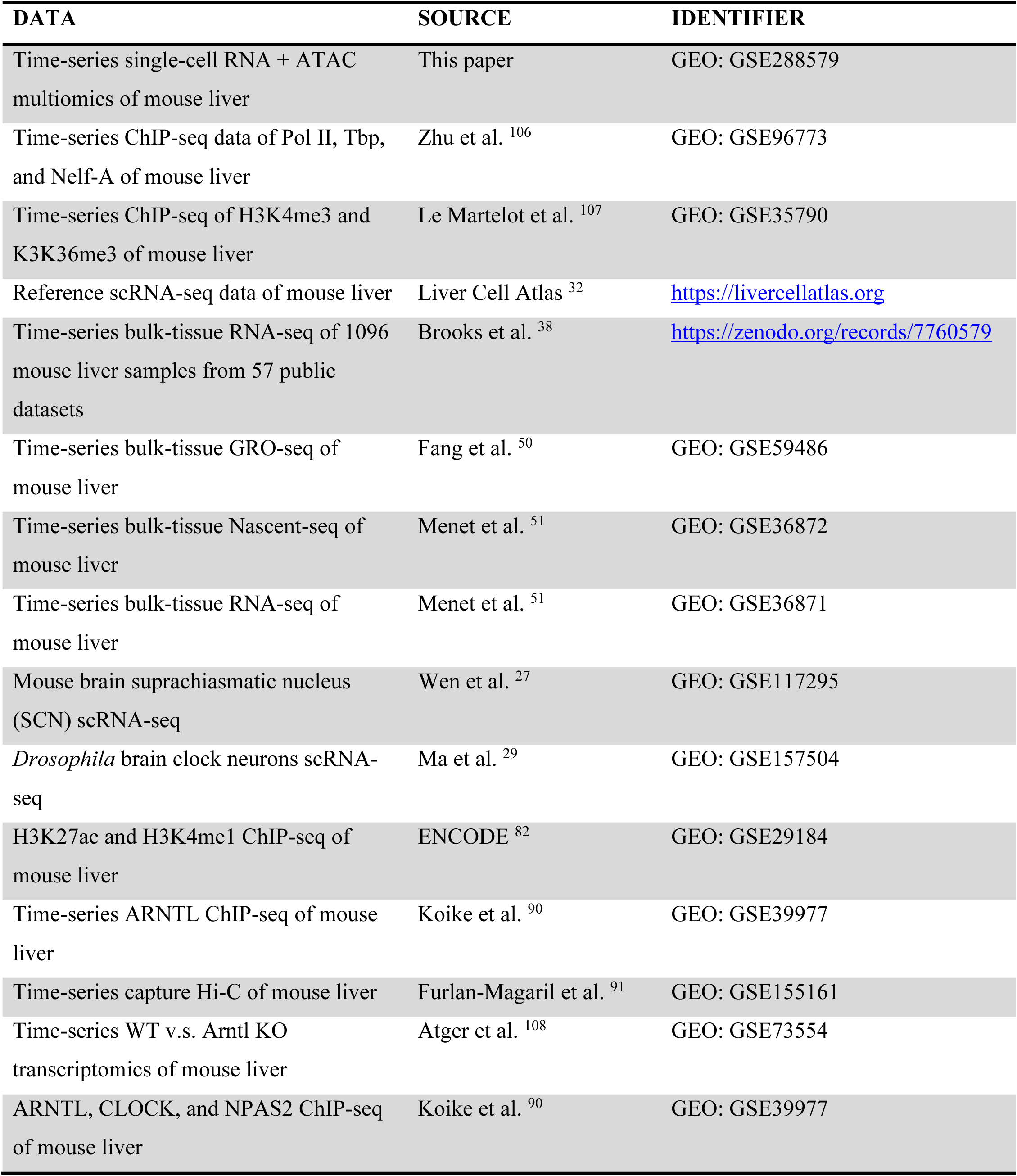

| DATA | SOURCE | IDENTIFIER |
| --- | --- | --- |
| Time-series single-cell RNA + ATAC multiomics of mouse liver | This paper | GEO: GSE288579 |
| Time-series ChIP-seq data of Pol II, Tbp, and Nelf-A of mouse liver | Zhu et al. <sup>106</sup> | GEO: GSE96773 |
| Time-series ChIP-seq of H3K4me3 and K3K36me3 of mouse liver | Le Martelot et al. <sup>107</sup> | GEO: GSE35790 |
| Reference scRNA-seq data of mouse liver | Liver Cell Atlas <sup>32</sup> | https://livercellatlas.org |
| Time-series bulk-tissue RNA-seq of 1096 mouse liver samples from 57 public datasets | Brooks et al. <sup>38</sup> | https://zenodo.org/records/7760579 |
| Time-series bulk-tissue GRO-seq of mouse liver | Fang et al. <sup>50</sup> | GEO: GSE59486 |
| Time-series bulk-tissue Nascent-seq of mouse liver | Menet et al. <sup>51</sup> | GEO: GSE36872 |
| Time-series bulk-tissue RNA-seq of mouse liver | Menet et al. <sup>51</sup> | GEO: GSE36871 |
| Mouse brain suprachiasmatic nucleus (SCN) scRNA-seq | Wen et al. <sup>27</sup> | GEO: GSE117295 |
| *Drosophila* brain clock neurons scRNA-seq | Ma et al. <sup>29</sup> | GEO: GSE157504 |
| H3K27ac and H3K4me1 ChIP-seq of mouse liver | ENCODE <sup>82</sup> | GEO: GSE29184 |
| Time-series ARNTL ChIP-seq of mouse liver | Koike et al. <sup>90</sup> | GEO: GSE39977 |
| Time-series capture Hi-C of mouse liver | Furlan-Magaril et al. <sup>91</sup> | GEO: GSE155161 |
| Time-series WT v.s. Arntl KO transcriptomics of mouse liver | Atger et al. <sup>108</sup> | GEO: GSE73554 |
| ARNTL, CLOCK, and NPAS2 ChIP-seq of mouse liver | Koike et al. <sup>90</sup> | GEO: GSE39977 |

## METHODS

### Mouse liver tissue collection

Experiments with mice were approved by the Texas A&M University Institutional Animal Care and Use Committee. Wild type adult males (C57BL/6NCrl strain; Charles River laboratories) were single housed under 12 h light:12 h dark and provided food and water *ad libitum*. Mice were euthanized by isoflurane anesthesia followed by decapitation, with four biological replicates combined at each time point for library preparation and sequencing. Livers were dissected quickly after euthanasia, briefly washed in ice-cold 1X PBS, snap-frozen in liquid nitrogen, and stored at −80°C until nuclei purification.

### Single-Cell RNA + ATAC multiomics

Nuclei isolation, library preparation, and sequencing of flash-frozen mouse liver tissue samples were performed at Singulomics Corporation (https://singulomics.com; Bronx, NY). Flash-frozen mouse liver tissue samples were processed to isolate nuclei using a combination of lysis, purification, and permeabilization steps. Nuclei were prepared by homogenizing the tissue in a lysis buffer, followed by filtration and resuspension in diluted nuclei buffer optimized for subsequent steps. This buffer formulation, containing magnesium and RNase inhibitors, ensured compatibility with downstream transposition and barcoding processes.

Purified nuclei were subjected to transposition in a reaction mix containing transposase, which preferentially fragmented open-chromatin regions and appended adapter sequences to the ends of DNA fragments. Polyadenylated RNA molecules within the nuclei were reverse-transcribed into full-length cDNA. These processes occurred within nanoliter-scale Gel bead-in-EMulsion (GEM), generated using the Chromium Next GEM platform. Each GEM contained the necessary reagents for DNA and RNA barcoding, ensuring unique molecular and nucleus-specific indexing.

Following GEM incubation, barcoded DNA and cDNA were recovered, purified, and amplified via PCR to generate sufficient material for library construction. ATAC libraries underwent size selection, while cDNA for gene expression libraries was enzymatically fragmented, end-repaired, and adapter-ligated prior to further amplification. Both libraries were subjected to QC to ensure high fidelity. Sequencing of the final libraries was performed on an Illumina NovaSeq platform using paired-end 150 bp reads (PE150), targeting a depth of ∼200 million reads per library.

### Demultiplexing, quantification, and quality control

Sequencing reads were analyzed with mouse reference genome mm10 using Cell Ranger ^109,110^ ARC 2.0.2. Aggregation of the samples was performed using the cellranger-arc aggr function, which subsampled from higher depth GEM wells until it equalized the median number of unique fragments per cell for each ATAC library and the mean number of reads mapped to the transcriptome per cell for each gene expression library. In addition to gene expression from the RNA assay and chromatin accessibility from the ATAC assay, we further utilized Signac ^105^ and chromVAR ^46^ to derive ATAC gene activity and motif score, respectively. Specifically, gene activity score ^45^ computes ATAC read counts per cell in the gene body and promoter region (2 Kb upstream of TSS), while motif deviation score ^46^ quantifies the overall accessibility within peaks sharing the same TF-binding motif while controlling for other technical biases.

We used scDblFinder ^111^ to remove cell doublets from both the RNA and ATAC modalities. We further adopted the following QC criteria and required each cell to have: (i) total number of reads between 1,000 and 100,000 for both RNA and ATAC, (ii) more than 500 detected features (genes and peaks), (iii) a TSS enrichment score between 3 and 20, (iv) a nucleosome signal below 2, and (v) mitochondrial read percentage under 5% (**Fig. S2**). These stringent thresholds ensure the inclusion of high-quality cells and features for downstream analysis. After QC, we retained a total of 19,273 cells with genome-wide measurements of 16,368 genes and gene activities, 149,302 peaks, and 746 motifs.

### Data normalization and integration

For data normalization, we applied different methods for each modality: RNA gene expression was normalized using sctransform, ATAC peak accessibility was normalized with Latent Semantic Indexing (LSI), and ATAC gene activity was normalized using logNormalize. Specifically, sctransform utilizes regularized negative binomial regression to normalize UMI count data. This approach effectively mitigates technical variability, such as differences in sequencing depth, while preserving biological heterogeneity. LSI applies term frequency-inverse document frequency (TF-IDF) for normalization, followed by singular value decomposition (SVD), correcting for differences in sequencing depth while retaining biologically meaningful variations. Log normalization of gene activity scores stabilizes variance, mitigates skewness, and enhances comparability across genes and cells. Motif deviation score is calculated by computing the difference between the observed number of fragments mapped to peaks containing the motif and the expected number, adjusting for technical confounders including GC content and the mean accessibility of the peak set ^46^. We used the the z-scores returned by chromVAR for each TF-binding motif in each cell.

Data integration was first performed across different time points within each modality for RNA gene expression, ATAC peak accessibility, ATAC gene activity, and motif deviation score, respectively (**Fig. S3**). Specifically, for each pair of datasets collected at two time points, we first identified “anchors”, consisting of two cells (with one cell from each time point) that were predicted to originate from a common biological state. An anchor score and weight were subsequently calculated, which were used for batch-effect correction using the technique adopted by the mutual nearest neighbor method ^112^. This procedure was iteratively performed to merge all six time points, ensuring that the integrated data accurately reflect biological differences while minimizing technical artifacts. We further carried out between-modality data integration to borrow information from different modalities using multiCCA ^47^ (**Fig. 1B**), a statistical method designed to identify shared patterns among multiple datasets. MultiCCA operates by finding linear combinations of variables within each dataset that are maximally correlated with those in other datasets, effectively capturing common sources of variation. This approach enables the alignment of distinct data types, such as RNA gene expression and ATAC peak accessibility.

### Annotating cell types, reconstructing metacells, and mapping spatial zonation

Cell-type annotations were transferred from the LCA ^32^, enabling the identification of hepatocytes, endothelial cells, fibroblasts, Kupffer cells, cholangiocytes, B cells, and T cells (**Fig. S4**). To ensure equal testing power across cell types that have different numbers of cells, we downsampled hepatocytes, endothelial cells, fibroblasts, and Kupffer cells to the same cell count for each cell type at each time point. We then performed cell-wise smoothing by pooling similar cells into metacells ^113^, representing distinct cell types and states. To determine the optimal resolution for metacell reconstruction, we used a list of 104 rhythmic and 113 arrhythmic genes ^53^ in mouse liver as positive and negative controls to generate the receiver operating characteristic (ROC) curve. The analysis indicated that a resolution of 2.5 yielded an area under the curve (AUC) greater than 0.8 for both RNA expression and ATAC activity (**Fig. S8**). The downsampling and cell aggregation procedures were repeated 20 times to ensure robustness and reproducibility of the results. To reconstruct the features for the metacells, we either took the average or calculated the fraction of cells with non-zero measures. Slingshot ^78^ was used to reconstruct the PN-to-CV trajectory and to infer pseudotime for each hepatocyte in the compartmentalized liver lobules. For simplified visualization, we further grouped the metacells into six spatial zones based on their inferred pseudotime.

### Circadian rhythm detection

To detect rhythmicity of RNA expression, ATAC activity and motif score, we applied both JTK_CYCLE ^54^ (implemented in MetaCycle ^55^) and harmonic regression ^56^. We combined the resulting nominal p-values using a heavy-tailed combination test ^57,114^ and adopted the Benjamini-Hochberg method for multiple testing correction (q-value < 0.01). We further applied a threshold on amplitude fold change for type I error control. The combination testing scheme integrates results from both parametric and nonparametric testing in a manner that goes beyond a simple intersection, and we refer readers to the statistics publication ^57^ for deeper theoretical aspects of this procedure. Phases were estimated through harmonic regression ^56^. Gene ontology enrichment analyses using cell-type-specific rhythmic genes were performed using clusterProfiler ^115^. We used PSEA ^61^ to identify phase-clustered functional enrichment using genes with cyclic RNA expression and CircaCompare ^63^ to test for significant phase shifting between different cyclic modalities. Motif enrichment analysis of rhythmic ATAC peaks was carried out using Signac ^105^ and JASPAR ^66^. Rhythmicity of bursting fraction and its comparison against that of mean expression in hepatocytes is calculated in a similar fashion.

### smFISH imaging, processing, and quantification

*Arntl* and *Per1* smFISH data were generated as previously described ^71^ on fresh-frozen liver cryosections (8 μm thick) embedded in O.C.T. Compound (Tissue-Tek; Sakura-Finetek), sampled every 3 hours from ZT00 to ZT21. RNAscope probes targeting *Arntl* (#438741) and *Per1* (#438751) were used according to the manufacturer’s protocol for the RNAscope Fluorescent Multiplex V1 Assay (ACDBio). Sections were counterstained with DAPI.

Imaging was conducted with a Leica DM5500 widefield microscope equipped with an oil-immersion ×63 objective lens. Z-stacks were acquired with 0.2 μm intervals. mRNA transcripts were quantified using ImageJ. Nuclei were detected through a series of image processing steps, including filtering to enhance features, thresholding to segment nuclei from the background, and watershed transformation to separate closely adjacent nuclei. PN and CV were manually annotated based on the presence or absence of the bile duct, as determined by DAPI staining. To detect fluorescent RNA-FISH spots, a Gaussian mixture model with two components was implemented to select only the larger dots colocalized with the genes in cell nuclei, ensuring accurate counting and an enhanced signal-to-noise ratio (**Fig. S28**). The total number of nuclei in each image was counted, and mRNA concentration was calculated as log₂ of the total number of nuclear smFISH transcripts divided by the total number of nuclei. The bursting fraction was calculated as the number of nuclei with active transcription divided by the total number of nuclei. Regions were selected for zoomed-in visualization at each ZT.

### Varying coefficient model for spatiotemporal analysis

We propose a statistical framework to test whether RNA expression (or ATAC activity) exhibits a significant spatial, circadian, or combined spatial and circadian pattern across spatial zonation and/or time points, focusing on one gene at a time. For cell *i*, let *y_i_* be its gene expression, *t_i_* be its ZT time point of collection, and *p_i_* be its pseudotime along the spatial gradient. We test one gene at a time and thus remove the gene index in the following models. Specifically, the null model *H*_0_ is specified as *y_i_* = *C* + *∈_i_*, where *C* denotes the baseline gene expression with error term *∈_i_* ∼ *N*(0, *σ*^2^). The circadian model *H*_1_ is specified as a cosinor curve ^116^ of form *y_i_* = *C* + *E* sin(*ωt_i_*) + *F* cos(*ωt_i_*) + *∈_i_*. For the spatial model *H*_2_, we carried out a basis expansion (polynomial of degree three by default) of the pseudotime to account for any nonlinear relationship *y_i_* = ∑*_k_ C_k_ϕ_k_*(*p_i_*) + *∈_i_*. Lastly, for the spatial and circadian model *H*_3_, we adopted a varying-coefficient model of form *y_i_* = ∑*_k_ C_k_ϕ_k_*(*p_i_*) + ∑*_k_ E_k_ϕ_k_*(*p_i_*) sin(*ωt_i_*) + ∑*_k_ F_k_ϕ_k_*(*p_i_*) cos(*ωt_i_*) + *∈_i_*, where the coefficients from the cosinor model are functions of the pseudotime upon basis expansions. We fit each of the four models and adopt likelihood ratio statistics for testing. Null log likelihood ratio statistics follows a chi-squared distribution; we perform a likelihood ratio test with finite-sample adjustments ^117^ to avoid inflated type I error.

### Linking spatiotemporal CREs to target genes

Hepatocytes are grouped by time points (ZT02, ZT06, ZT10, ZT14, ZT18, ZT22) or zonation (zones 1 to 6) and randomly sampled without replacement to reconstruct four replicates per time point or zone, ensuring equal testing powers. We calculate the mean RNA expression and mean ATAC peak accessibility for each group and scan for significant association between a target gene and a peak region located within 500 Kb of the gene’s TSS via a correlation-based test. A rhythmic CRE is linked to a circadian gene across time points with a significant correlation coefficient greater than 0.5. Similarly, a spatial CRE is linked to a spatial gene across spatial zonation with a significant correlation coefficient greater than 0.5. A spatial and rhythmic CRE is linked to a spatial and circadian gene with correlation coefficients significantly exceeding 0.5 under both contexts.

### CRE-TF-gene regulatory trios in circadian control

Metacell-specific omics profiles were used as input for TRIPOD ^44^, a nonparametric computational framework designed to analyze single-cell multiomic data and identify regulatory relationships between CREs, TFs, and target genes. The candidate CRE regions are defined as 200 Kb upstream and downstream of the TSSs. An q-value threshold of 0.01 was adopted to assess significance. The results from TRIPOD were further validated through chromatin-state annotation, promoter capture Hi-C, and TF ChIP-seq, demonstrating the robustness of the findings.

## DATA AND CODE AVAILABILITY

Raw and processed single-cell RNA and ATAC multiomic data from this study have been deposited at GEO (https://www.ncbi.nlm.nih.gov/geo) under the accession number GSE288579. All original code is publicly available at the GitHub repository https://github.com/yuchaojiang/circadian_sc_multiomics. An R shiny app is available at https://mouselivermultiomics.shinyapps.io/singulomics_rshiny.

