## Supplementary Figures for "Single-Cell Multiomic Analysis of Circadian Rhythmicity in Mouse Liver"

**Fig. S1.** Data processing and analytical framework, including bioinformatic pre-processing, quality control, both within- and between-modality data integration, and outlines of downstream analyses.

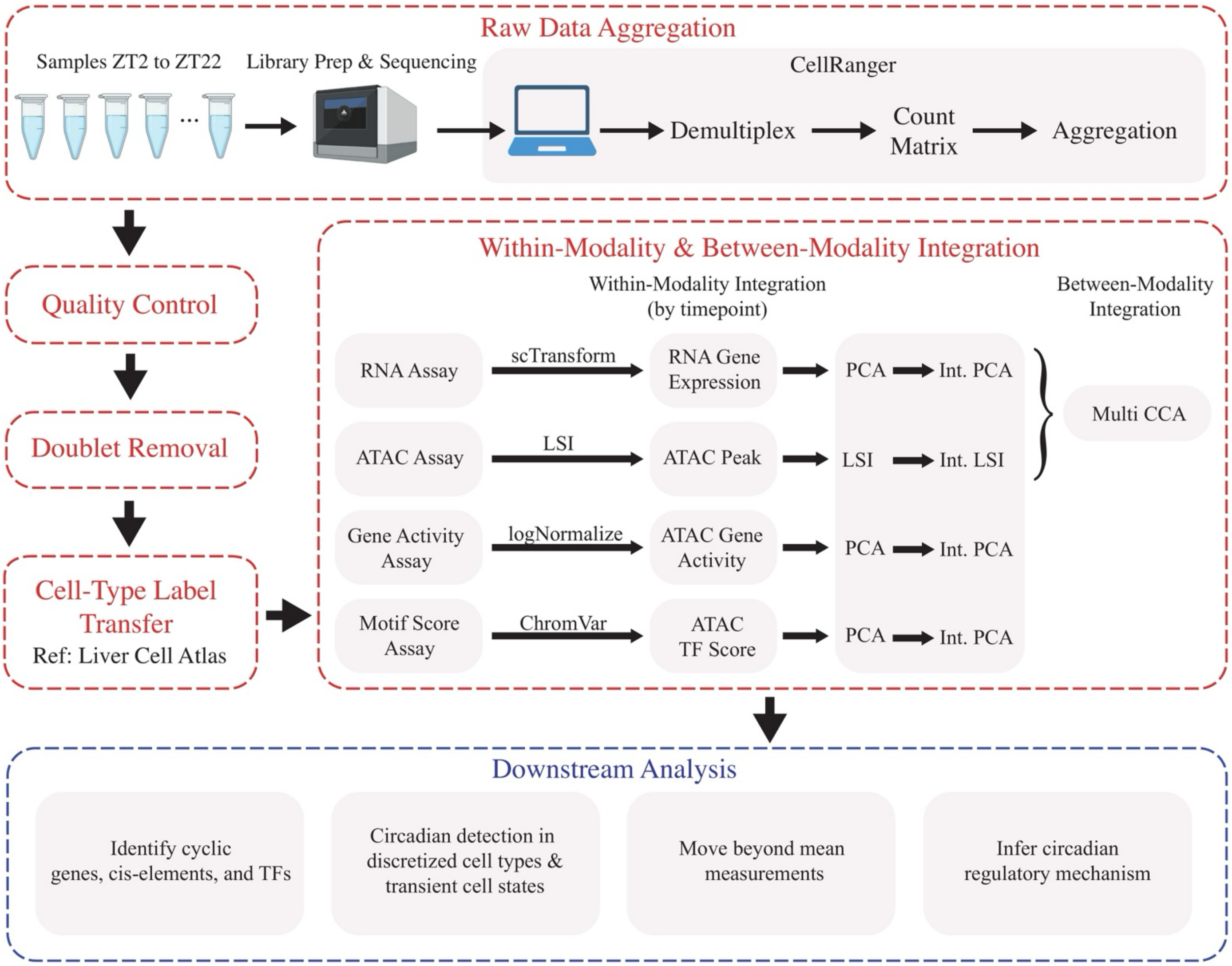

**Fig. S2.** Quality control (QC) metrics: (A) before and (B) after QC. Specific metrics include: number of ATAC counts (nCount_ATAC), number of ATAC peaks (nFeature_ATAC), transcriptional start site (TSS) enrichment score based on the ratio of fragments centered at the TSS to fragments in TSS-flanking regions (TSS.enrichment), nucleosome banding pattern quantified by the approximate ratio of mononucleosomal to nucleosome-free fragments (nucleosome_signal), number of RNA counts (nCount_RNA), number of genes (nFeature_RNA), and percentage of reads that map to the mitochondrial genome (percent.mt).

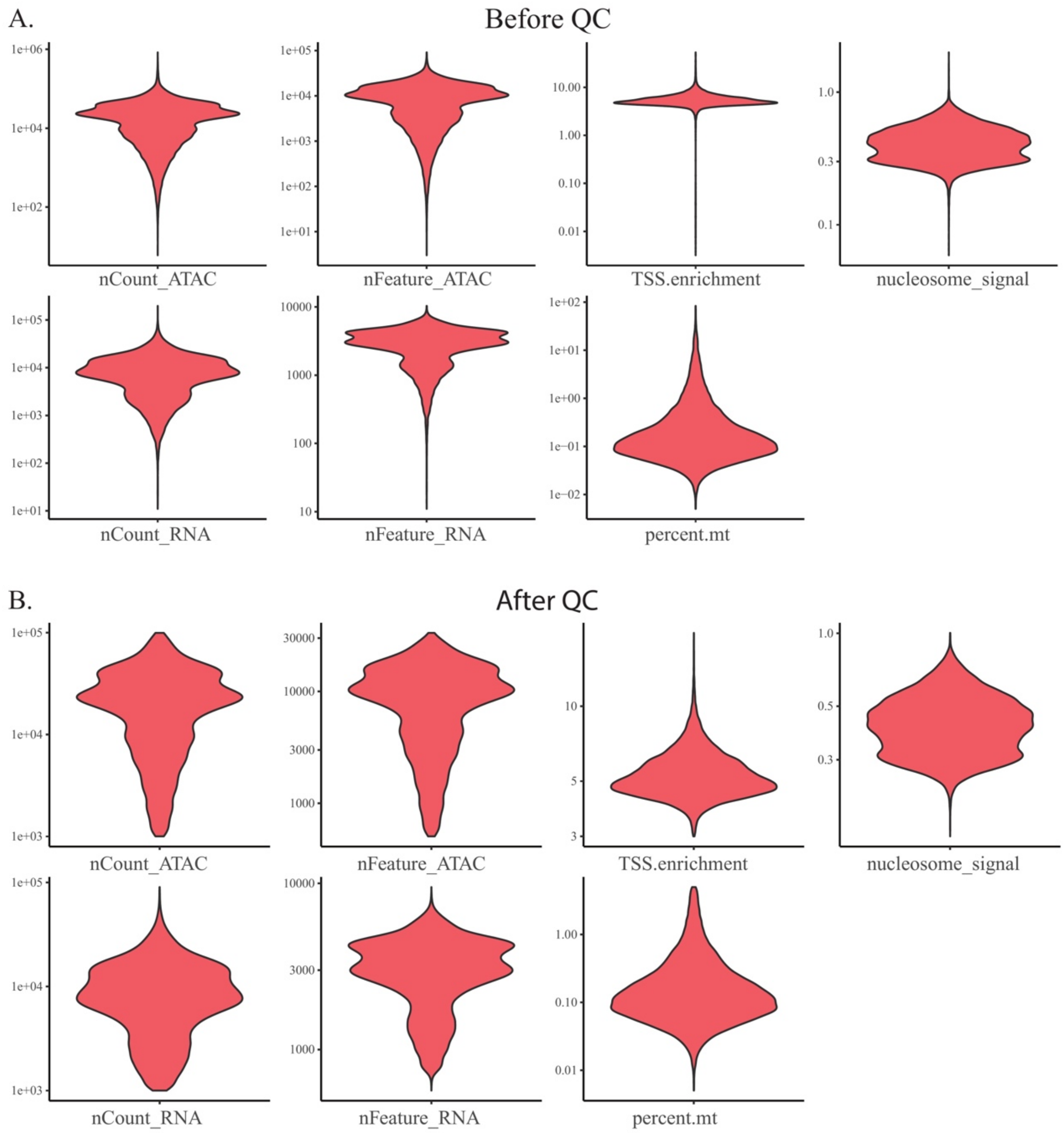

**Fig. S3.** Within-modality data integration across circadian time points. (A) Before and (B) after data integration for RNA, ATAC peak, ATAC activity, and MOTIF scores. Both scRNA-seq data and scATAC-seq data show strong batch effect before correction, each sample form distinct clusters. After batch-effect correction, samples from different circadian time points are integrated into cohesive clusters.

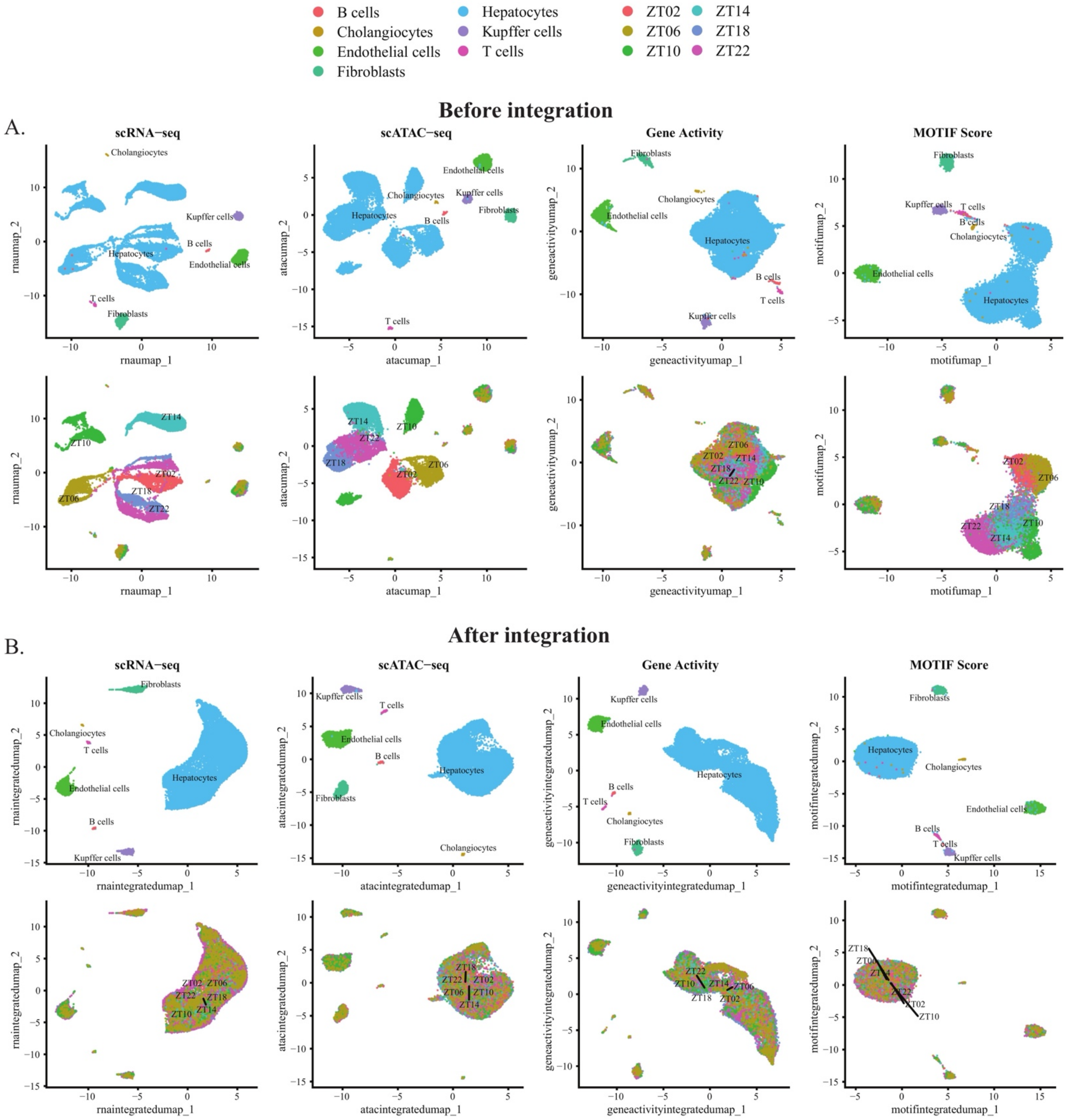

**Fig. S4.** Cell-type label transfer. (A) Single-nucleus RNA sequencing data generated by nucSeq from the Liver Cell Atlas (LCA) [1] used as reference. (B) Number of cells at different time points in different annotated cell types. (C) Prediction scores (between zero and one and akin to probabilities of assignment) for cell-type label transfer from the LCA.

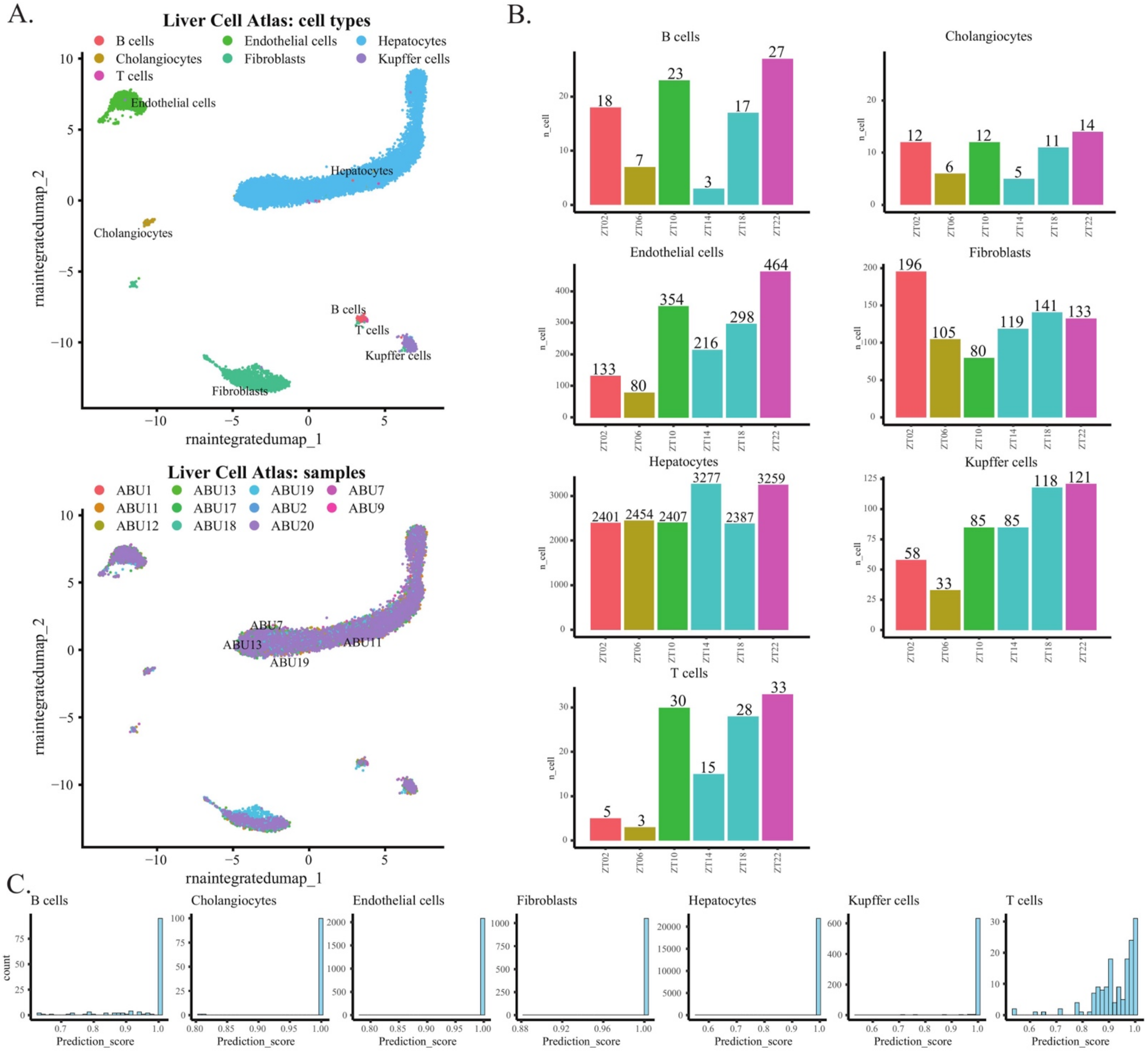

**Fig. S5.** Cell-type-specific marker genes show consistency between our data and the LCA [1]. (A) Heatmap of top cell-type-specific gene markers. (B) Cell-type-specific marker genes from the LCA are highly correlated with our data. (C) Heatmaps depict the relative gene expression of cell-type-specific DEGs reported from the LCA.

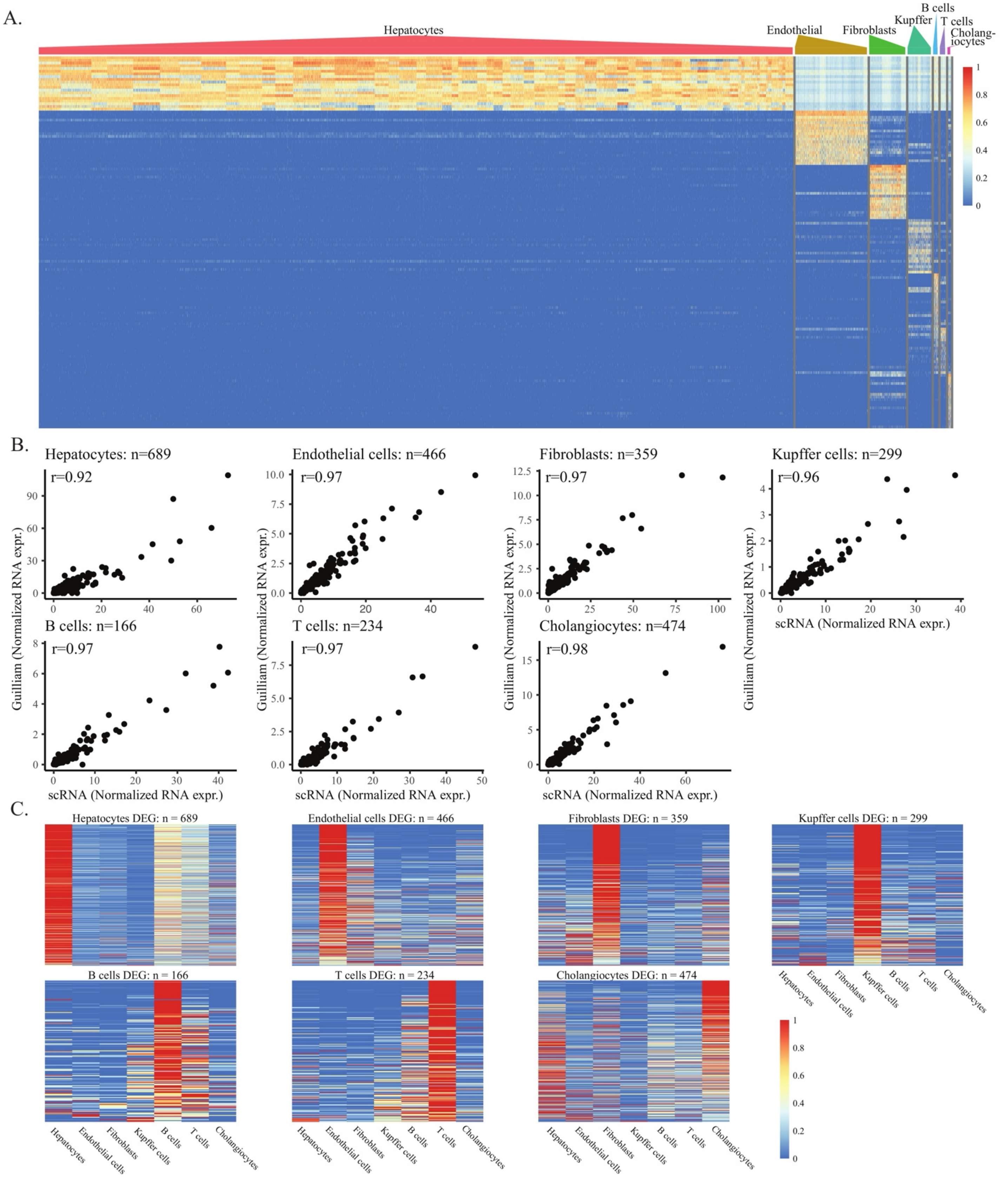

**Fig. S6.** High-quality ATAC data identified candidate cis-regulatory elements (CREs). (A) ATAC peaks significantly overlap with previously reported DNase I hypersensitive sites (DHSs) in mouse liver [2], demonstrating good ATAC quality. (B) Boxplot shows the identified and overlapped peaks are significantly enriched with H3K27ac and H3K4me1 histone markers than randomly sampled peaks. (C) Coverage plots depict that our ATAC peak are concordant with mouse DHSs for representative circadian genes (*Arntl*, *Nr1d1*, and *Dbp*).

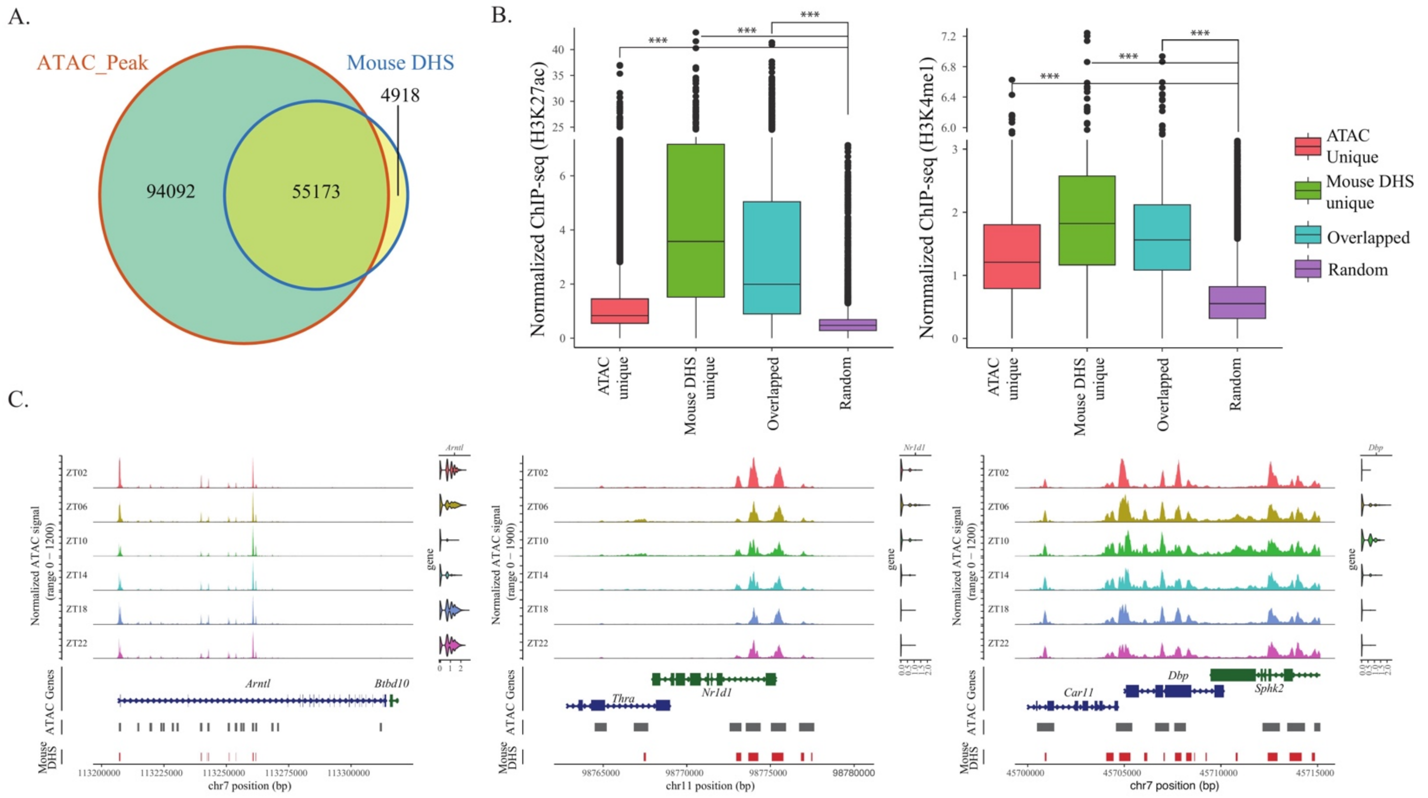

**Fig. S7.** Prediction of ZT time points using bulk data to demonstrate utility and good RNA expression data quality. (A) Architecture of deep neural network for ZT prediction. (B) Prediction performance on bulk-tissue GRO-seq [3], Nascent-seq [4], and RNA-seq [4] data of mouse liver. (C) 1096 time-series RNA-seq mouse liver samples from 57 public repositories [5] were obtained and analyzed; only the control groups of each study were included, to create comparable data.

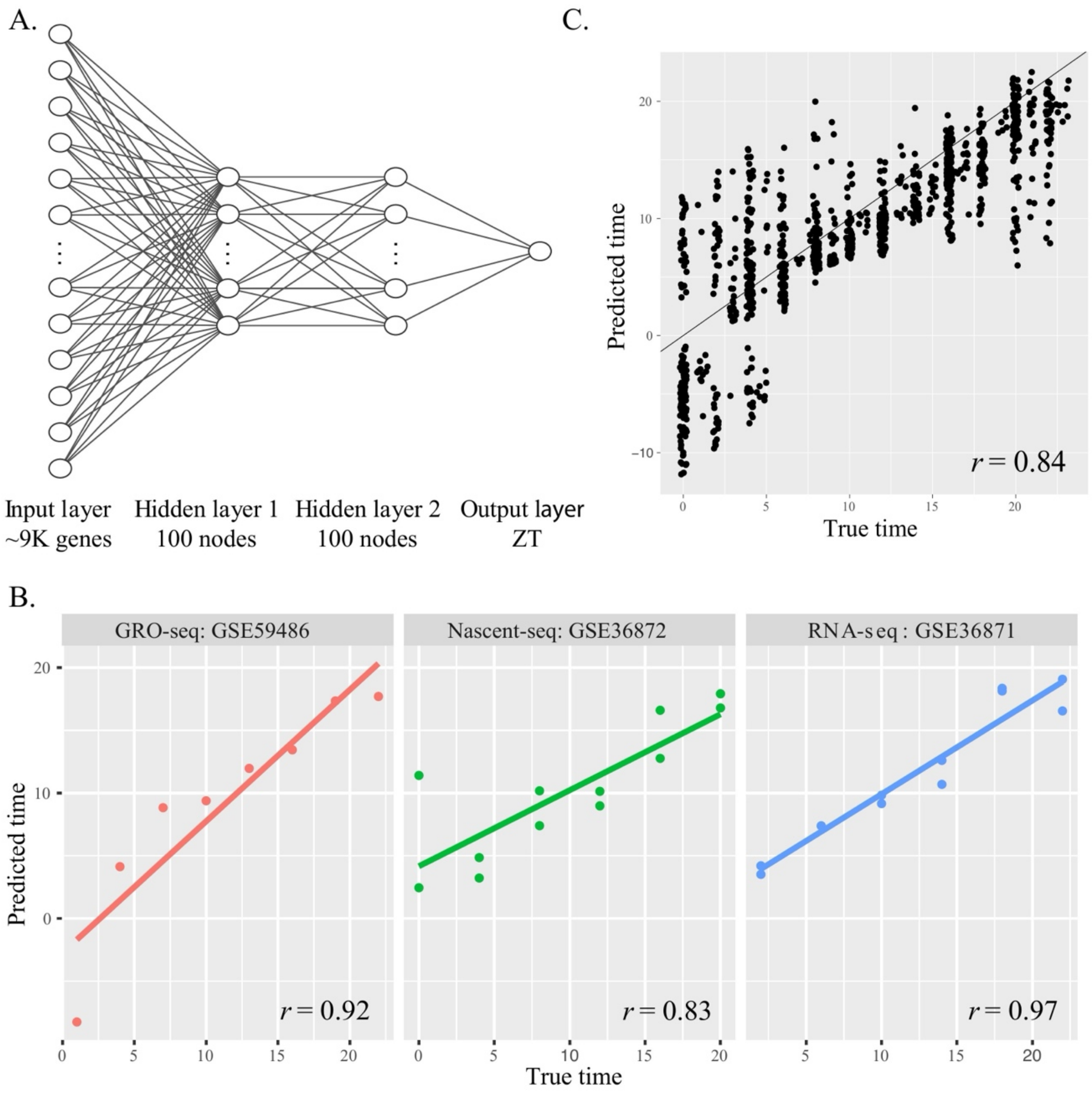

**Fig. S8.** Resolution tuning for the construction of metacells of hepatocytes. (A) Dot plot of resolution vs n_meatacell. (B) ROC curve of RNA assay and ATAC gene activity assay in various resolutions. (C) AUC with varying number of metacells. n_metacell=25 and resolution=2.5 were chosen, with AUC higher than 0.8 for both RNA and ATAC. (D) Number of cyclic genes (p.adj < 0.01) with varying number of metacells. (E) Density plot of adjusted p values in different resolutions. (F) Construction of metacells along the central vein (CV) to portal node (PN) gradient. (G) Cells collected at different ZT time points have the same pseudotime distribution (i.e., at each segment along the transient gradient, circadian time points are well mixed).

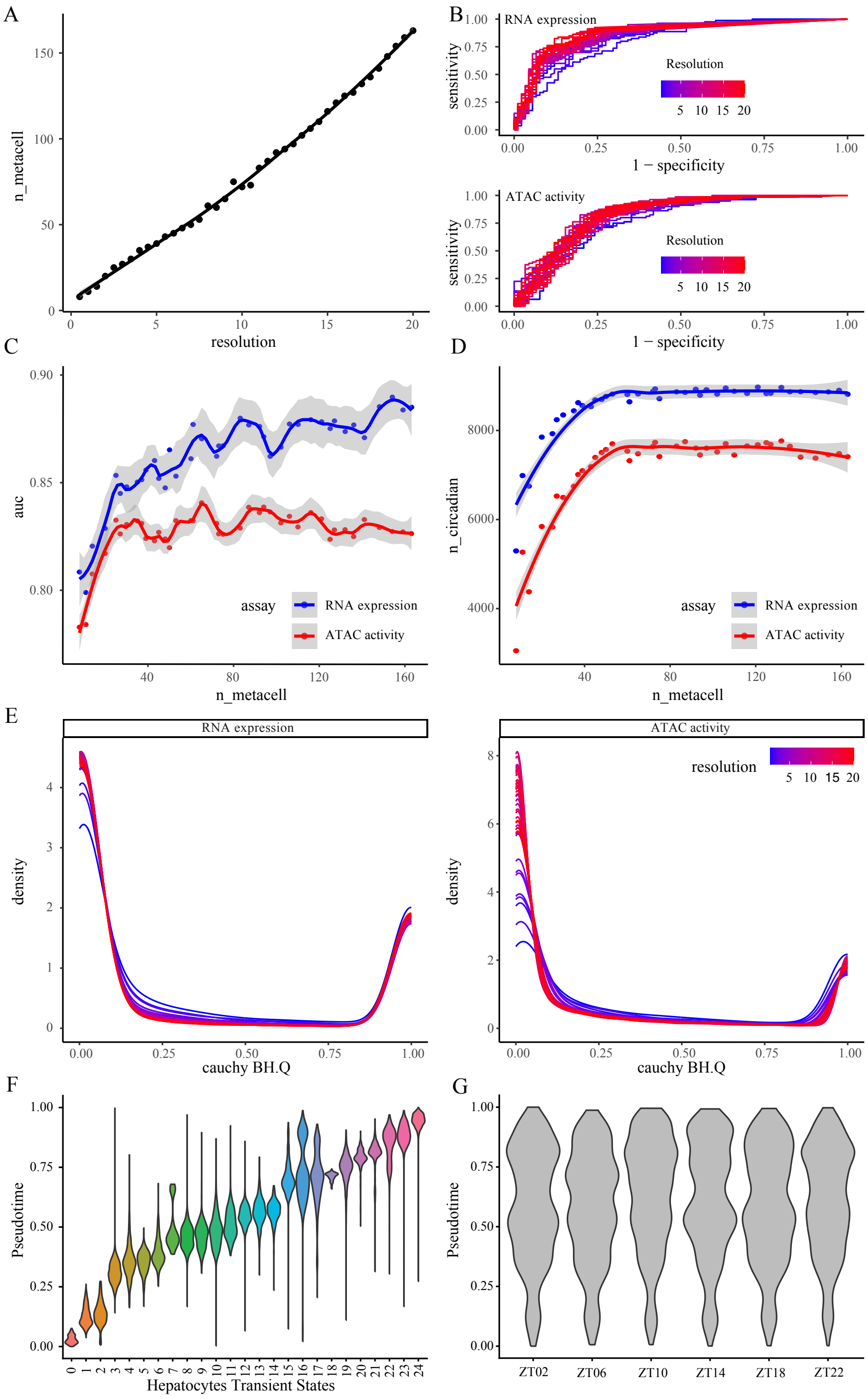

**Fig. S9.** Detecting circadian rhythmicity by JTK Cycle and harmonic regression, followed by a heavy-tailed combination test. (A) Methods for detecting circadian rhythmicity and estimating phase.(B) P-value correlation between JTK Cycle and harmonic regression (HR). (C) Distribution of nominal p-value from permutation to generate true null for harmonic regression (HR), JTK Cycle and JTK Cycle + HR Cauchy combination.

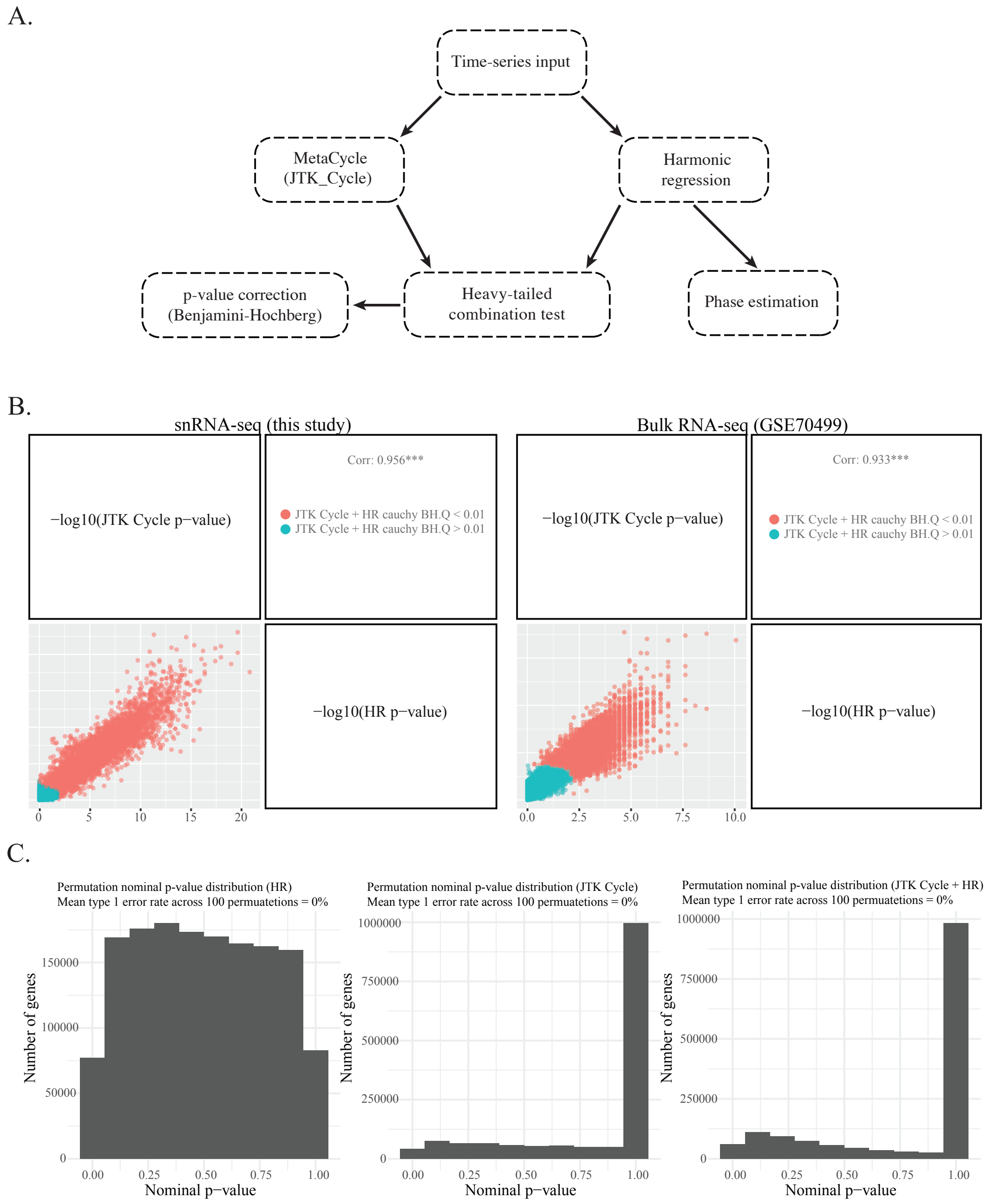

**Fig. S10.** Cell-type-specific rhythmic RNA expression and ATAC accessibility for core clock genes with error estimates and testing results.

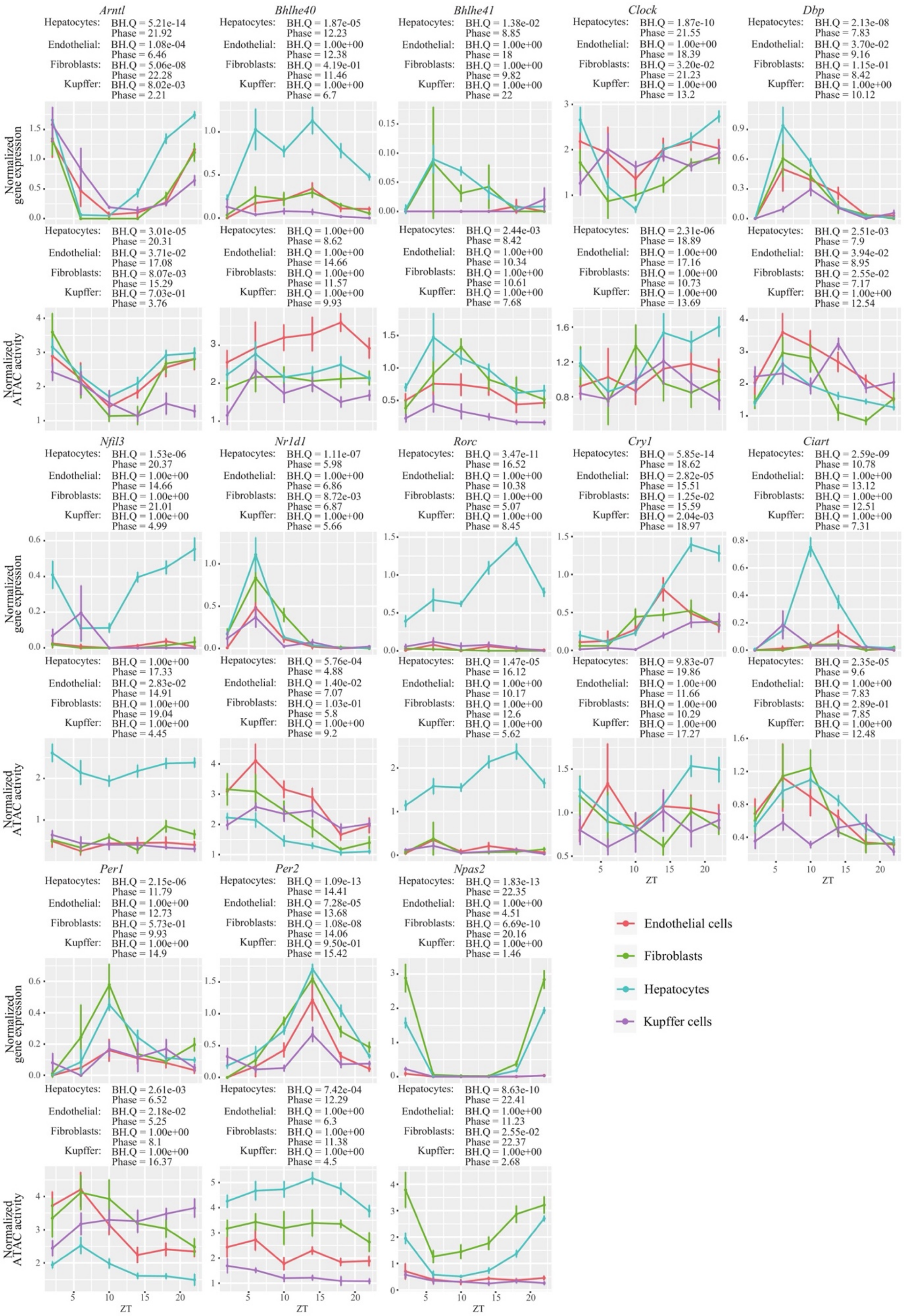

**Fig. S11.** Hepatocytes harbor substantially higher numbers of read counts (nCount_RNA) and gene counts (nFeature_RNA) in (A) this study and (B) the LCA [1]. This seemingly high difference is biological – cell size has been shown to correlate with the total number of transcripts and thus the total number of reads per cell [6], and hepatocytes are markedly larger than other liver cell types [7]. Importantly, to make it comparable across cell types, the raw read counts were normalized by the library size (i.e., the total number of reads, which is also highly correlated with the number of genes detected).

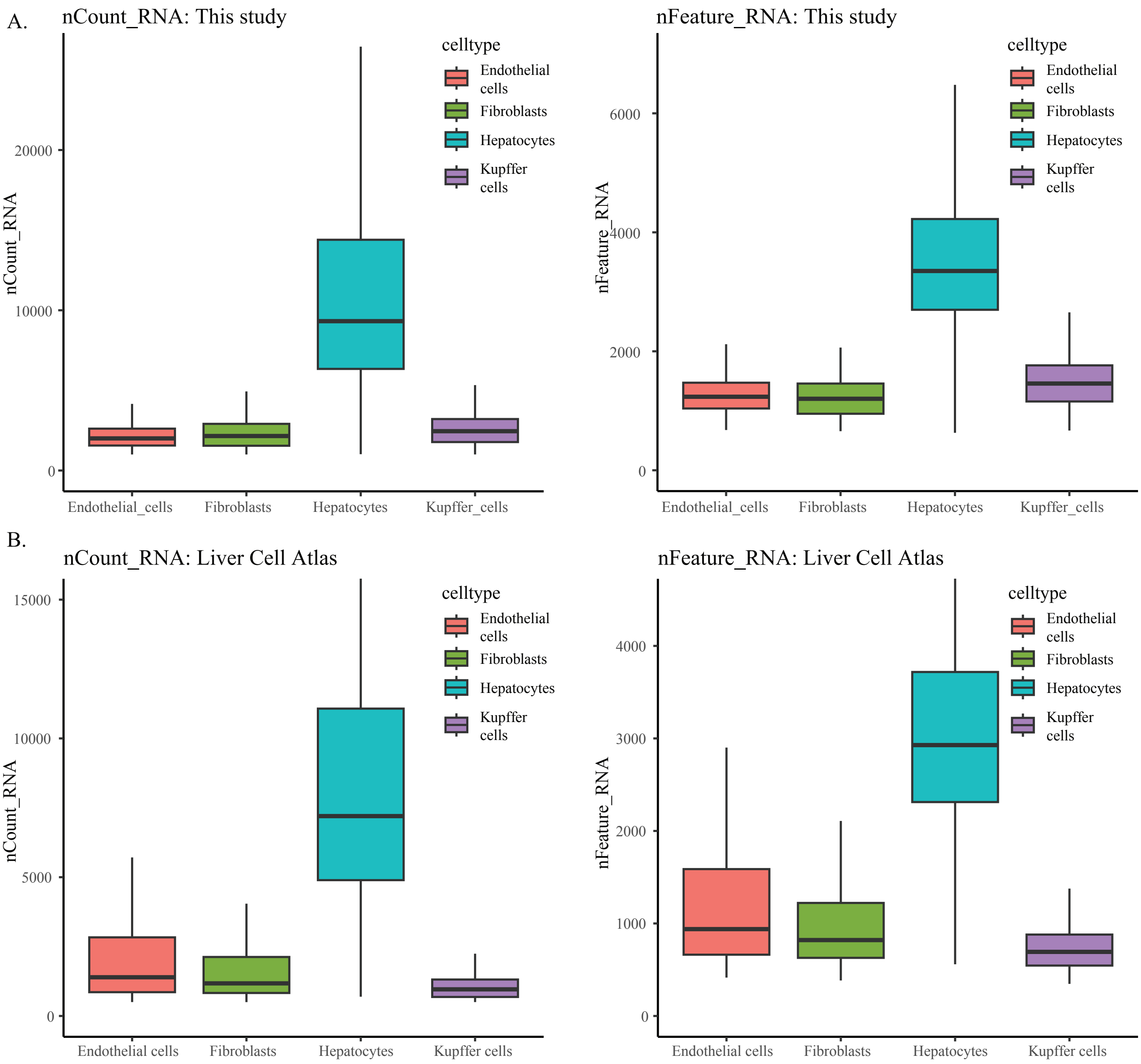

**Fig. S12.** Cell-type specific phase set enrichment analysis (PSEA). (A) Hepatocytes, (B) Endothelial cells, (C) Fibroblasts and (D) Kupffer cells.

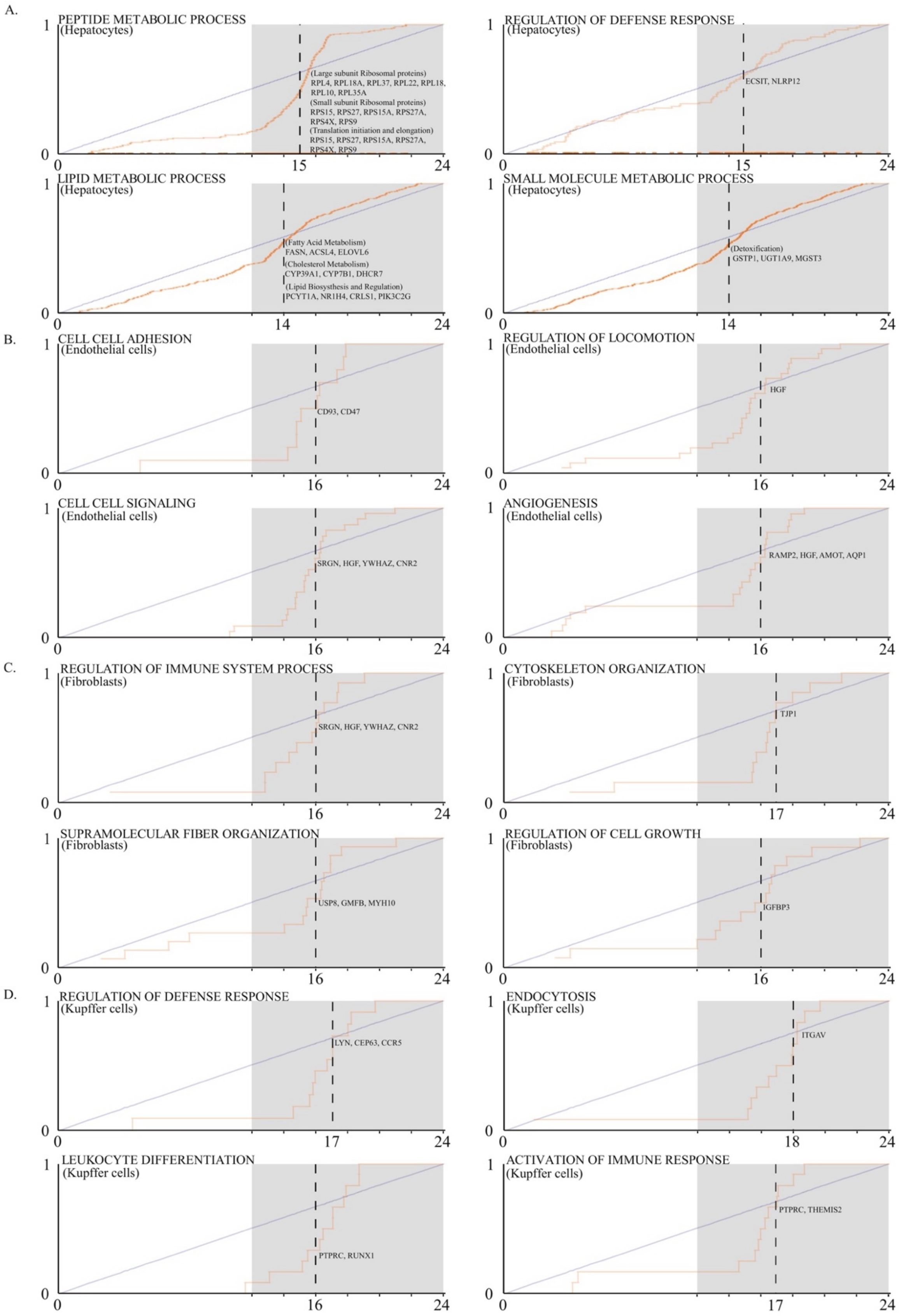

**Fig. S13.** Phase shift analysis of RNA expression and ATAC activity. (A) Phases of RNA expression and ATAC activity are highly correlated. We tested for significant phase shifts between the two modalities using CircaCompare [8]. (B) Genes with significantly smaller ATAC phases compared to RNA are longer and exhibit larger relative amplitude (rAMP).

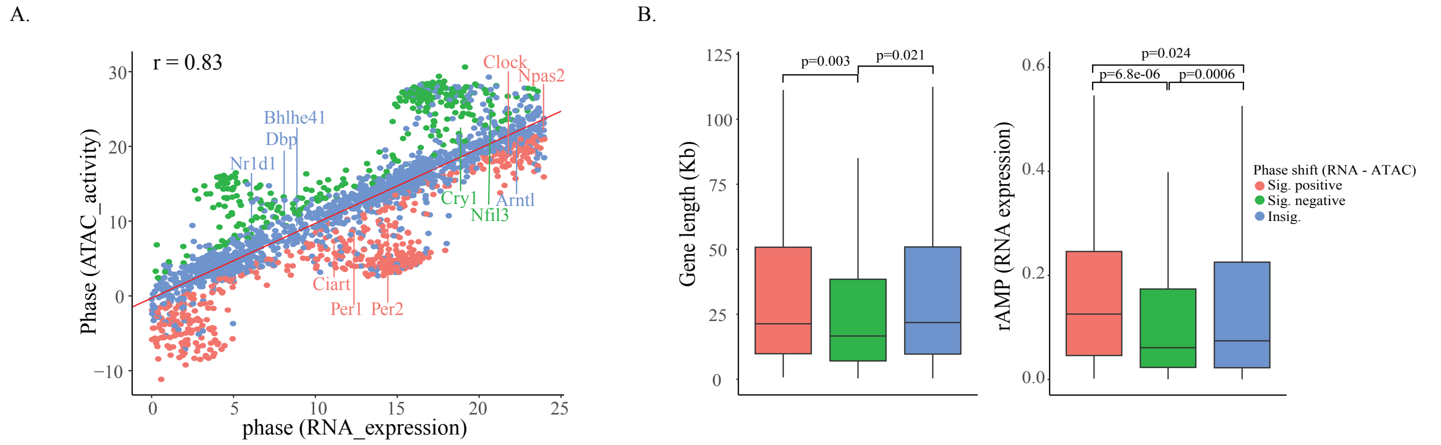

**Fig. S14.** TF-specific testing of circadian rhythm. (A) Overlap of rhythmic TF motif score, TF gene RNA expression, and TF gene ATAC activity. (B) Cell-type-specific rhythmic TF expression, activity, and motif deviation.

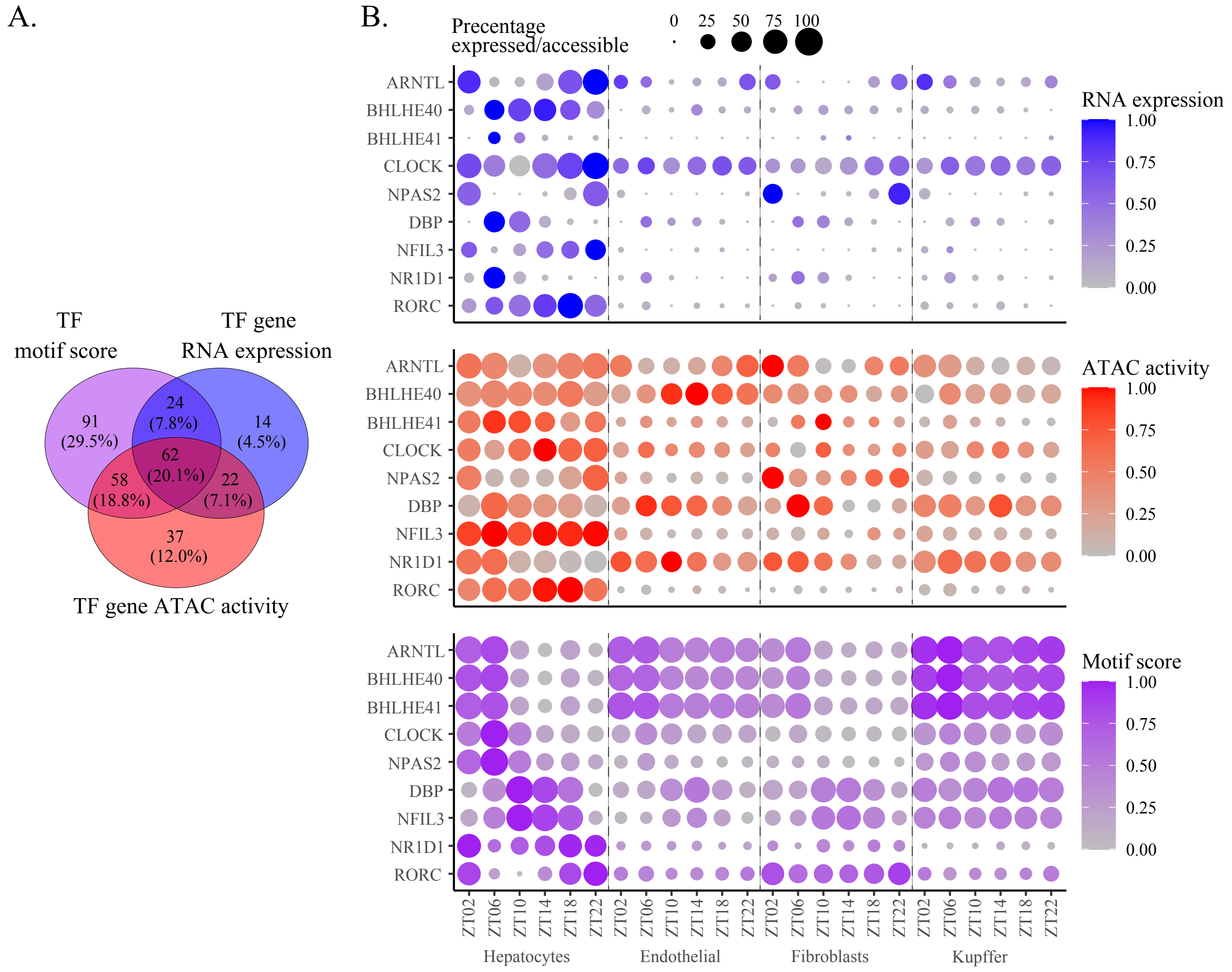

**Fig. S15.** Circadian rhythmicity regulated by burst fraction. Results are adapted from Phillips et al. [9] (https://github.com/naef-lab/CircadianSMFISH). (A) *Nr1d1*, *Bmal1*, and *Cry1* smFISH of mouse fibroblasts. Estimates of (B) burst frequency and (C) burst size of the three rhythmic genes. Rhythmic mean expression can be recapitulated by rhythmic burst frequency, while burst size is invariant to circadian time but only proportional to cell size.

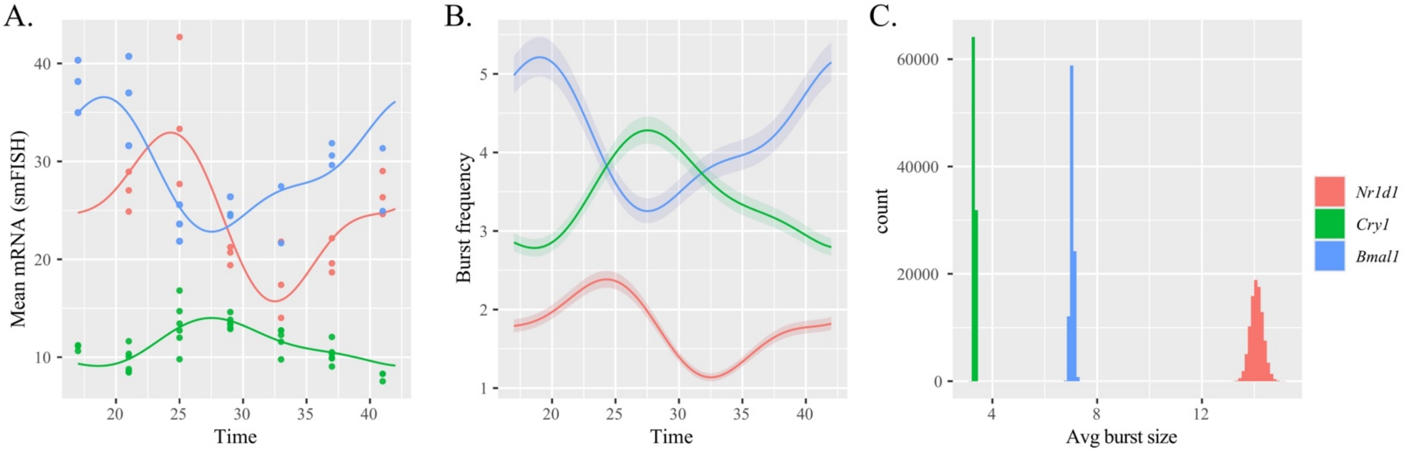

**Fig. S16.** Circadian rhythmicity detected by different summary statistics from cell-level read counts. (A) Upset plot of cyclic genes detected by different summary statistics. On the global scale, non-zero mean (surrogate for burst size) returns the least number of rhythmic genes, and when it does, it captures the highly/constitutively expressed genes that can generally be detected by the other measures. (B) Smooth scatter plots and Spearman correlation coefficients between each pair of summary statistics. Non-zero proportion is nearly perfectly correlated with Gini index and coefficient of variation. Furthermore, non-zero proportion, compared to non-zero mean, has a much higher correlation coefficient with mean. Results are shown across the top 2000 cyclic genes in the genome.

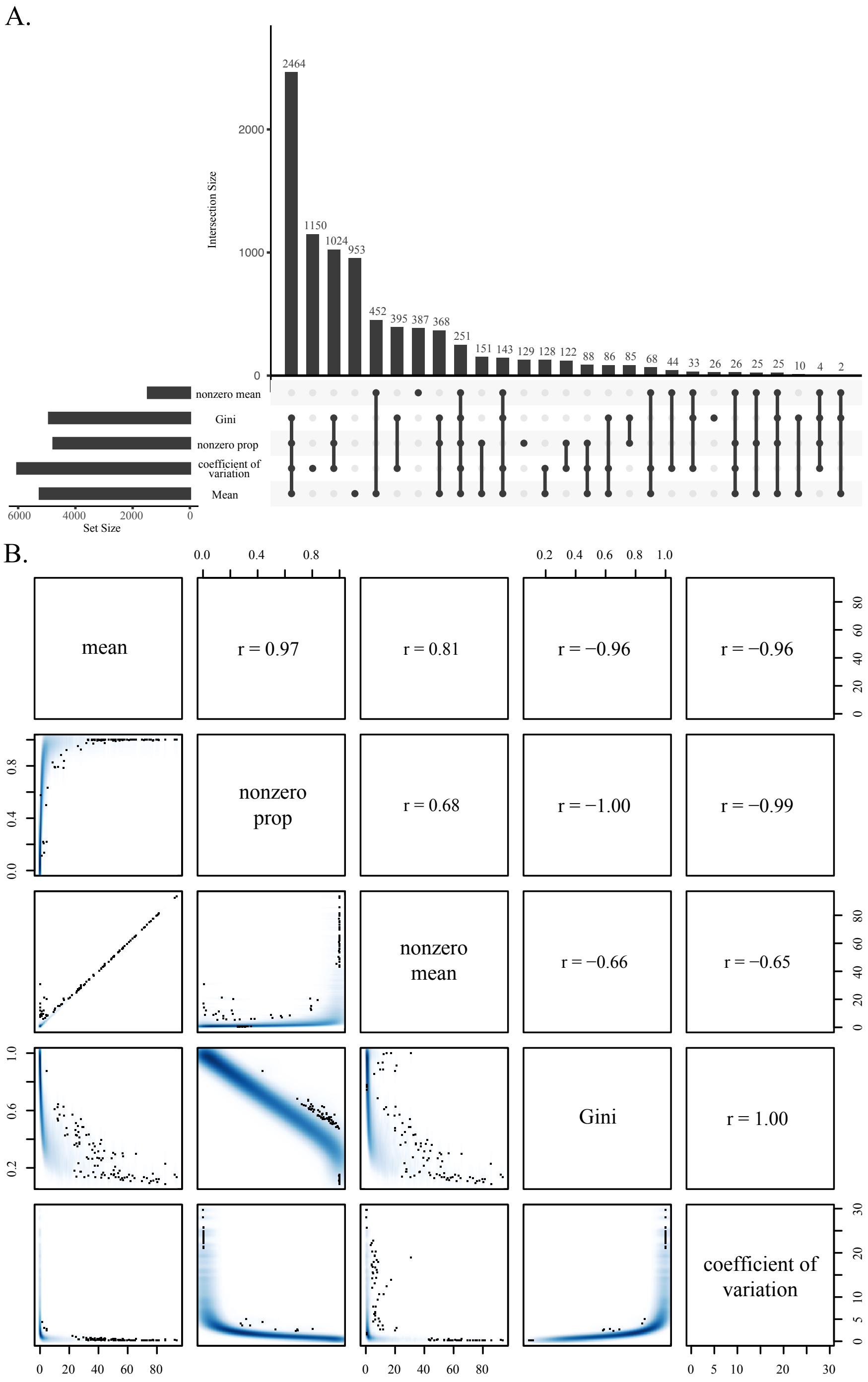

**Fig. S17.** Mouse SCN scRNA-seq data from Wen et al. [10]. (A) Dimension reduction of mouse SCN. (B) Mean expression and non-zero proportion of core clock genes in Neurons1. (C) Venn diagram (mean v.s. non-zero proportion). (D) Boxplot demonstrate that genes identified by mean rhythmicity have higher average expression and lower bursting fraction.

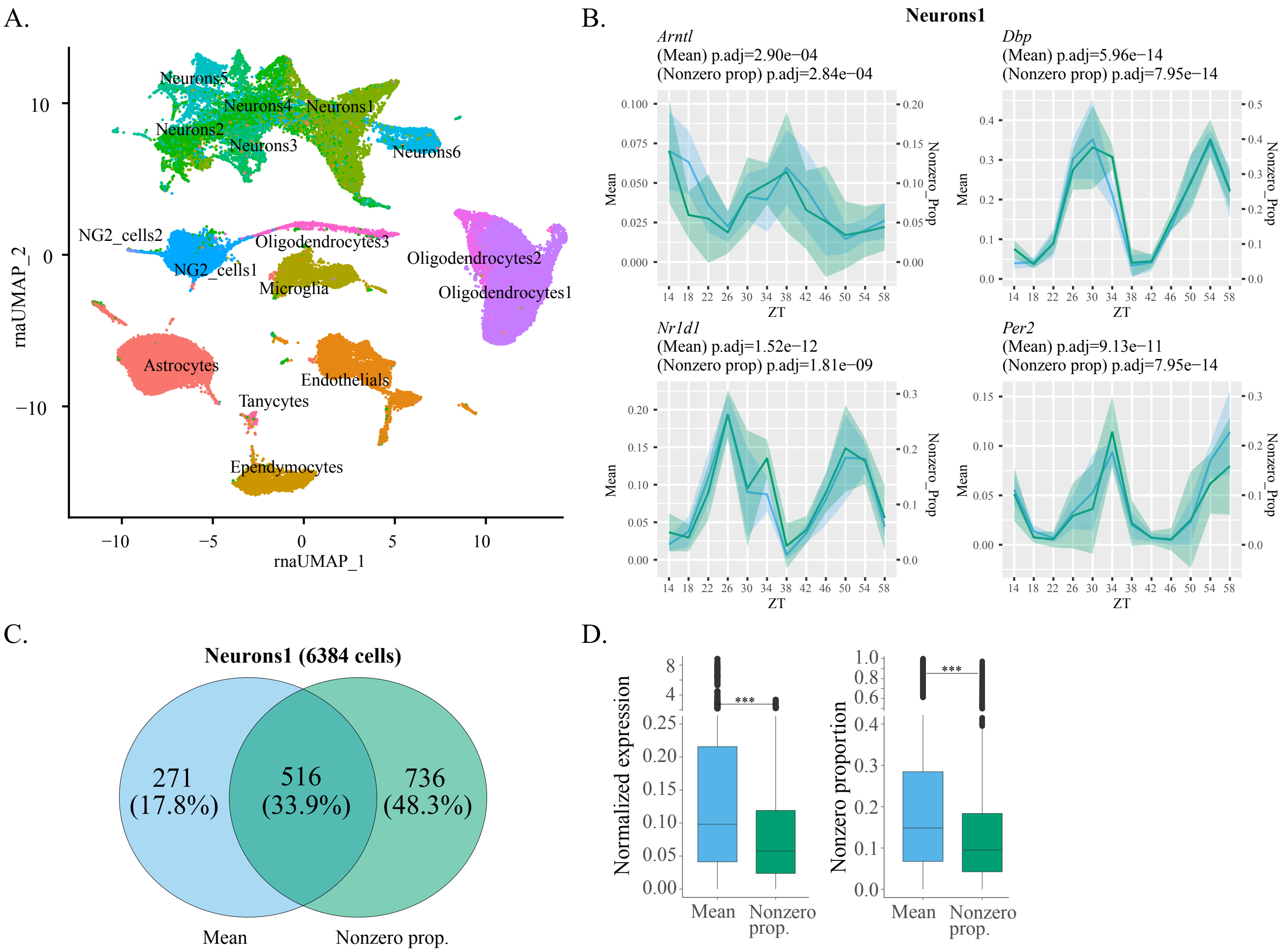

**Fig. S18.** Drosophila neuron scRNA-seq data from Ma et al. [11]. (A) Dimension reduction of 17 cyclic neuronal clusters (with significant rhythmicity for both *tim* and *Clk*) identified by the original study. (B) Mean expression and non-zero proportion of core clock genes *Tim* and *Clk*. (C) Venn diagram (mean v.s. non-zero proportion). (D) Boxplot demonstrate that genes identified by mean rhythmicity have higher average expression and lower bursting fraction.

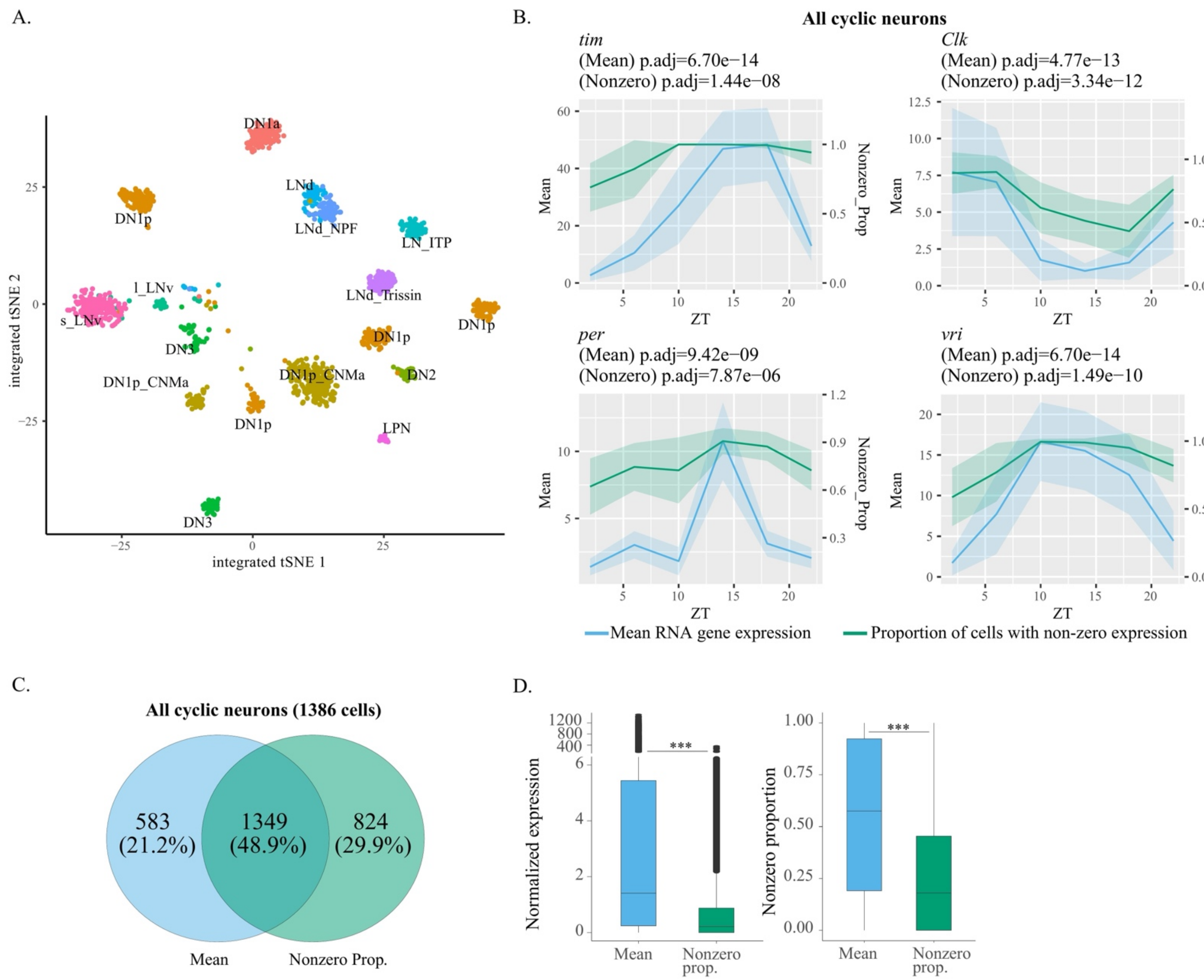

**Fig. S19.** Visualization of circadian genes, spatial genes, and spatial & circadian genes.

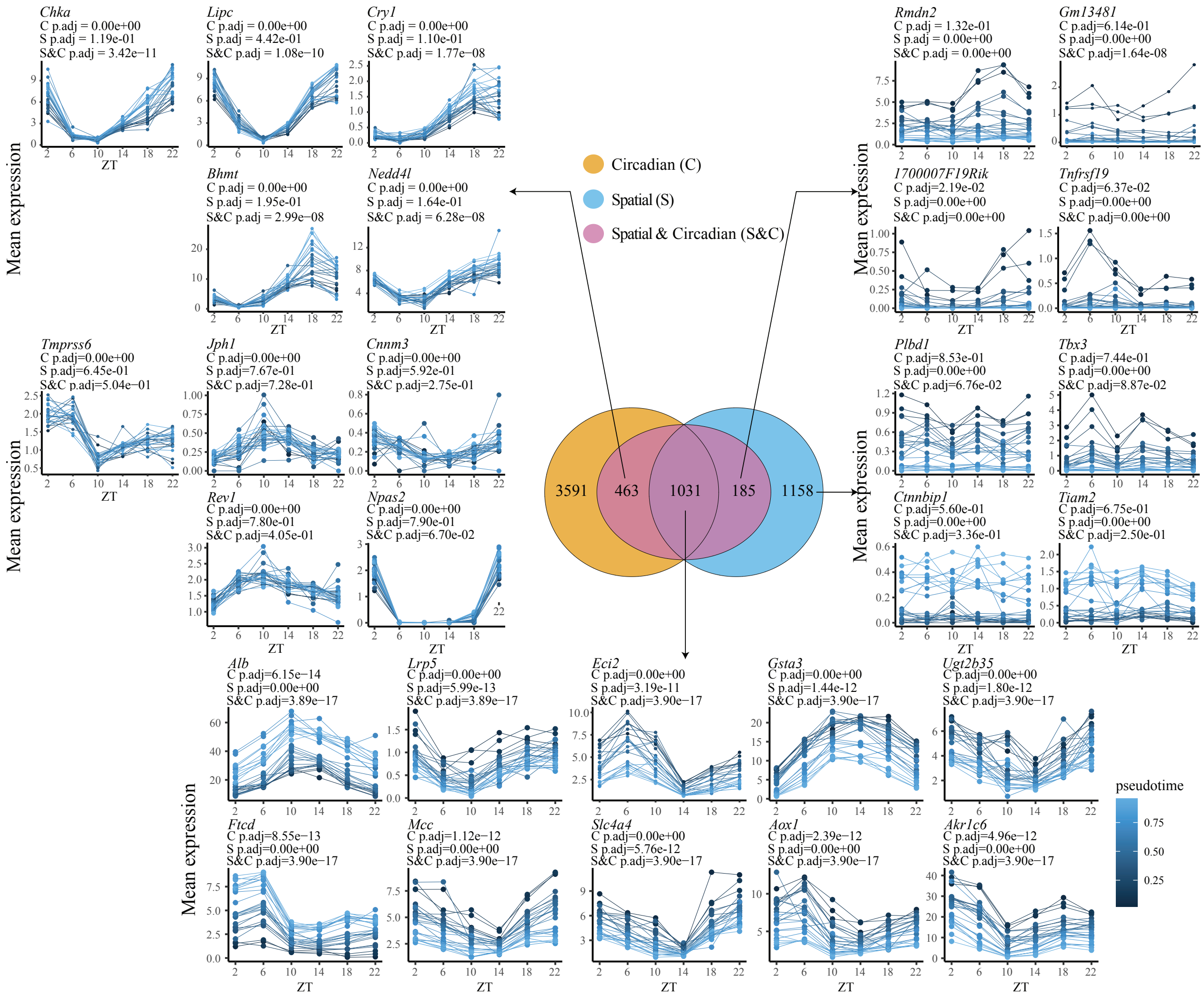

**Fig. S20.** Spatial and circadian testing results of clock reference genes.

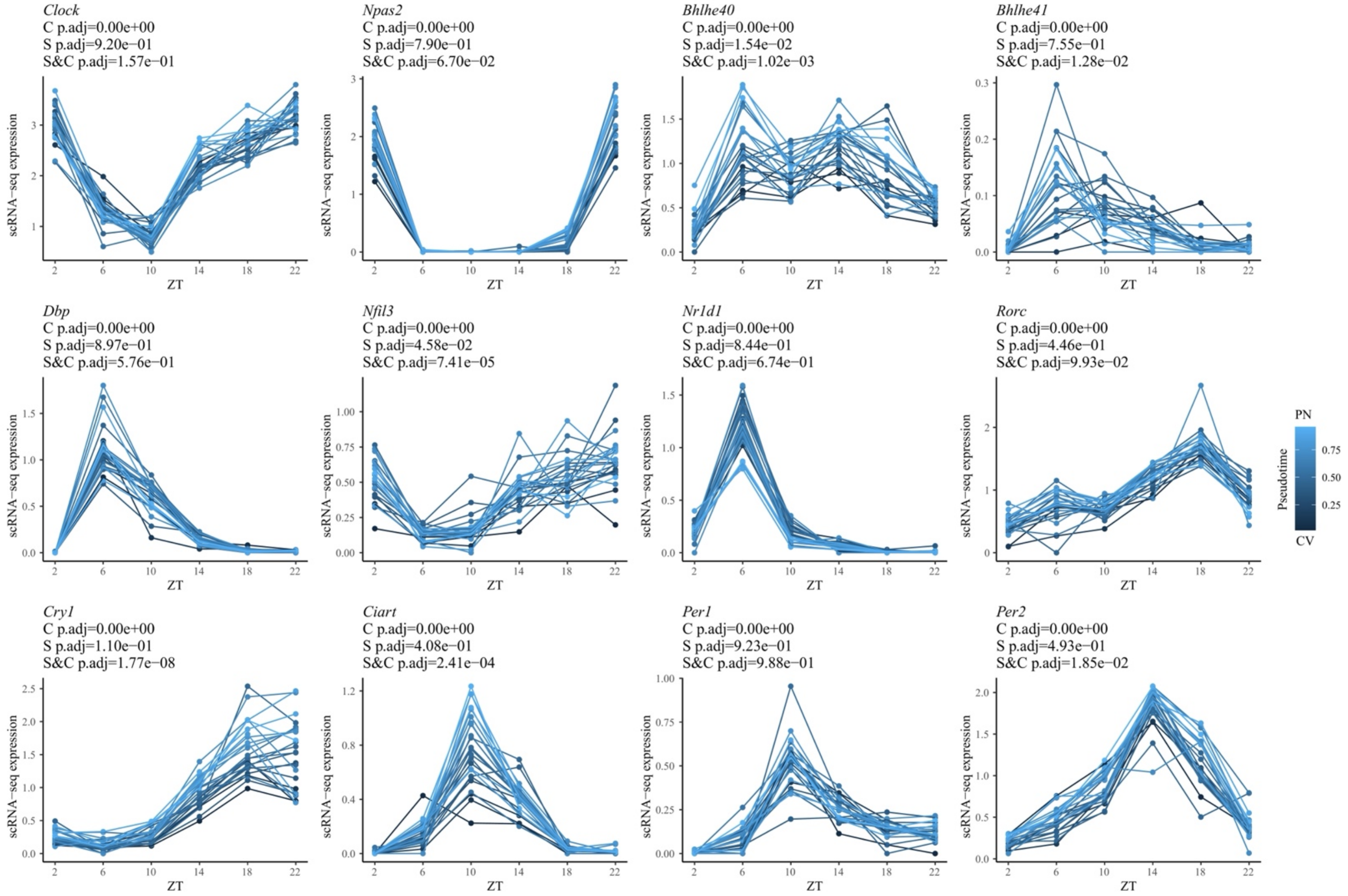

**Fig. S21.** Transient genes enriched near the portal node (PN) and central vein (CV). (A) RNA expression and ATAC activity heatmaps of 1,031 spatial and circadian genes from CV to PN. (B) KEGG pathway enrichment analysis of the 1,031 spatial and circadian genes that are enriched in either CV or PN. We find that CV-enriched genes are involved in pathways such as peroxisome [12], PPAR signaling pathway [13], drug metabolism [14], and bile secretion [13], in concordance with previous conclusions. PN-enriched genes, on the other hand, are involved in pathways such as complement and coagulation cascades [15].

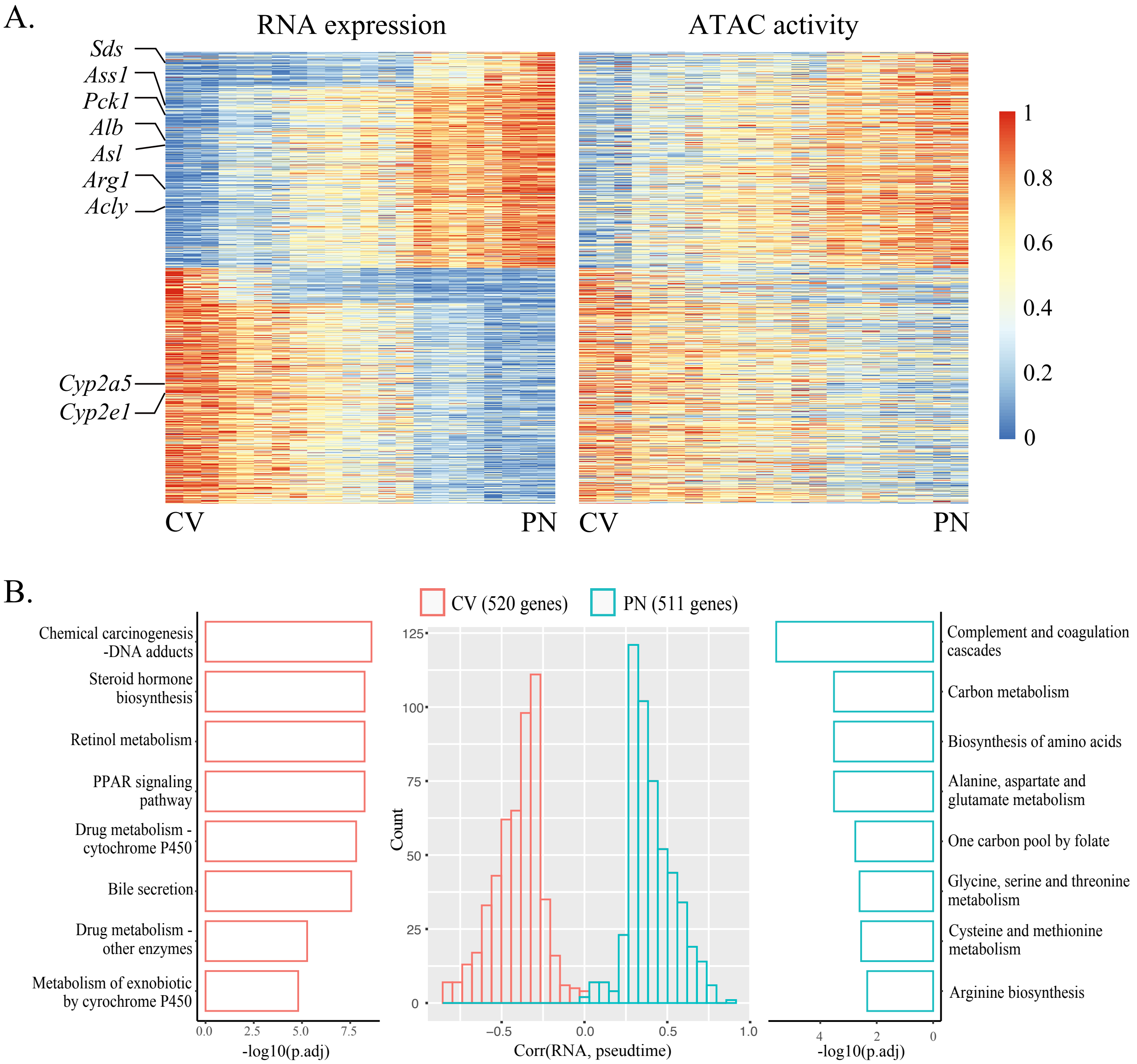

**Fig. S22.** Circadian, spatial, and spatial and circadian ATAC gene activities exhibit similar patterns as RNA gene expressions.

A.

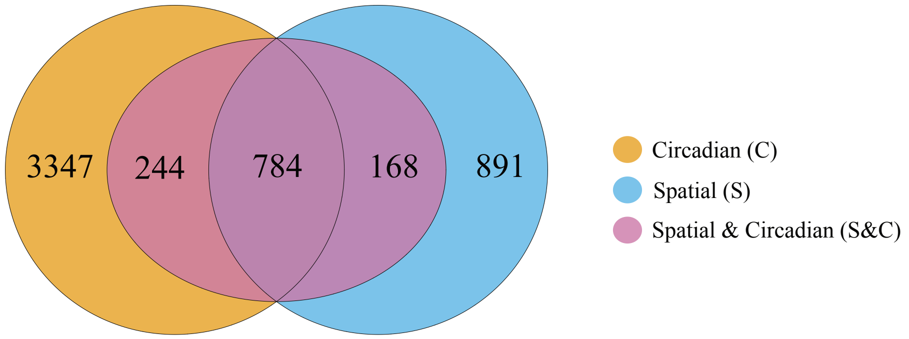


B.

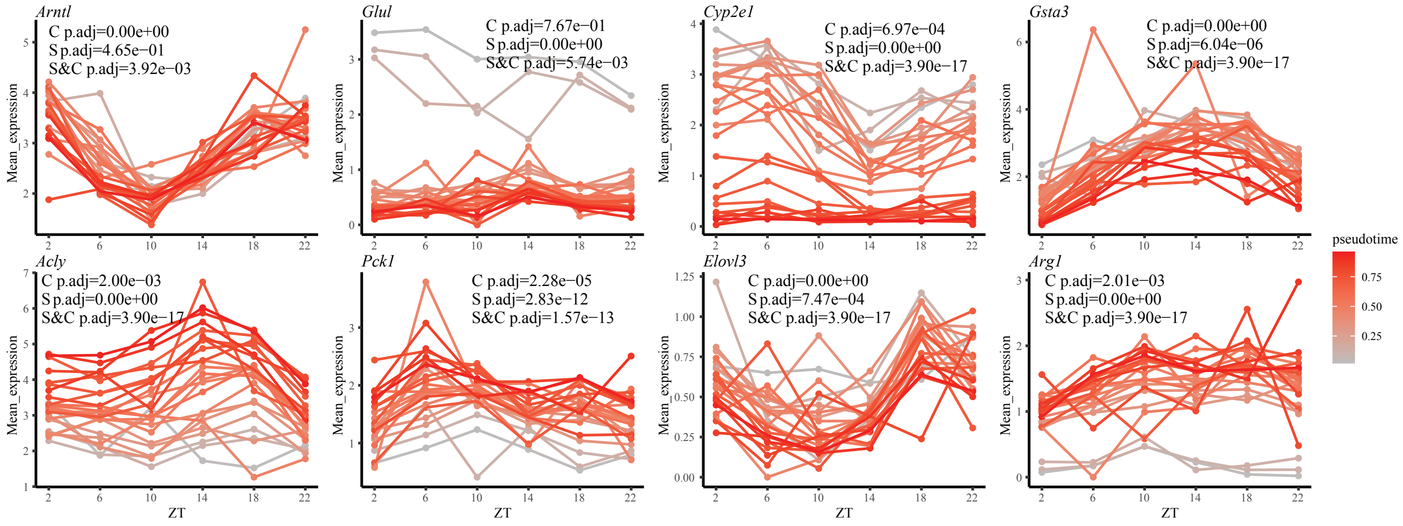

**Fig. S23.** Linked ATAC peaks temporally correlated with *Clock*, *Cry1*, *Dbp*, *Nr1d1*, and *Rorc*.

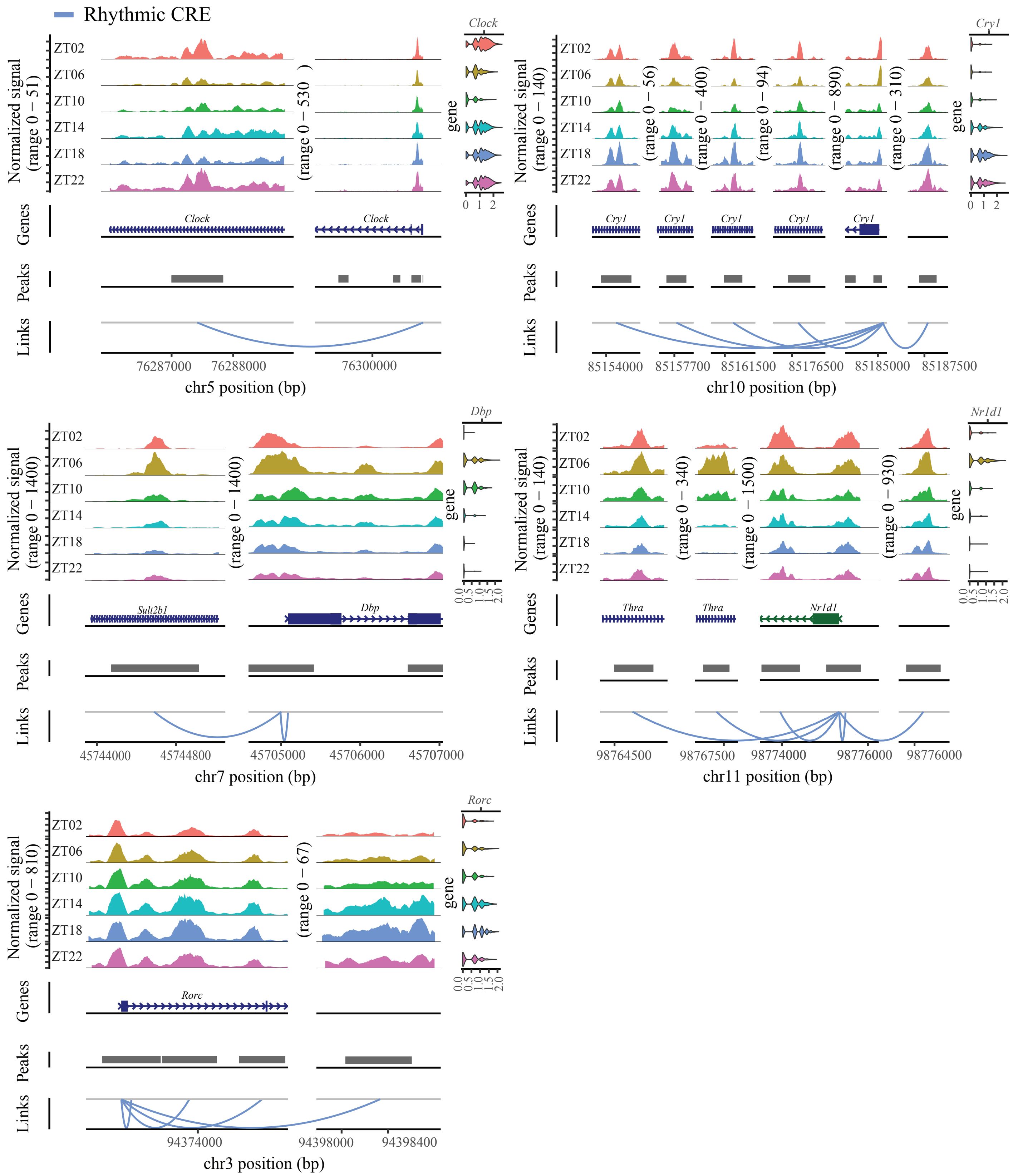

**Fig. 24.** Top significantly enriched motifs from spatial, rhythmic, and spatial and rhythmic CREs.

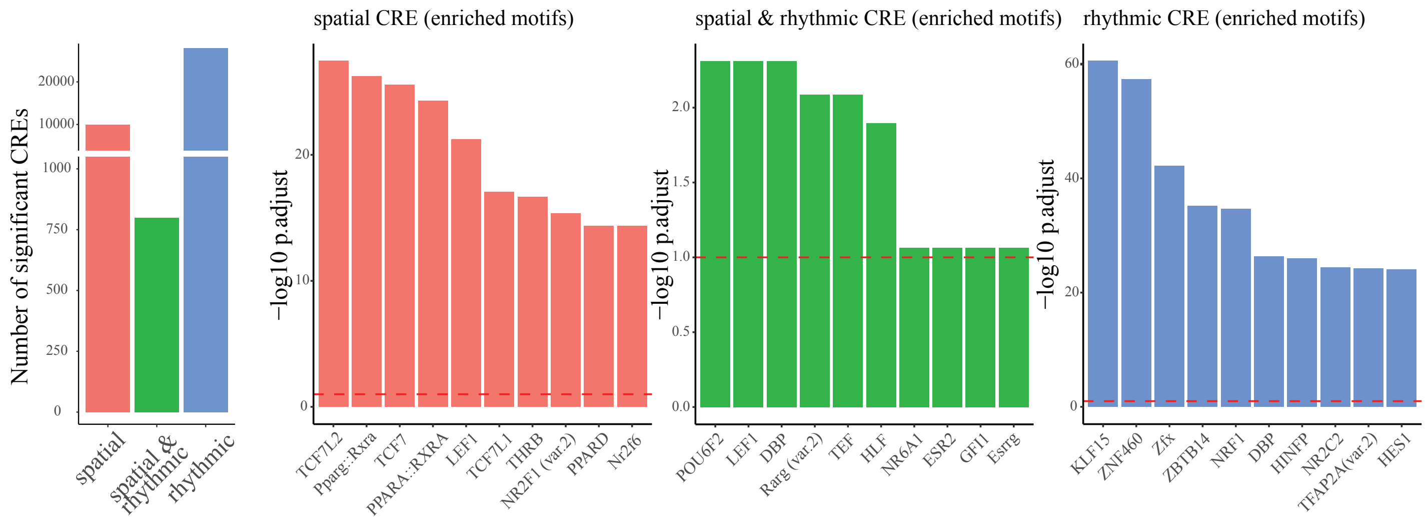

**Fig. S25.** Predicted CRE-TF-gene trio relationships with *Arntl* as the target gene.

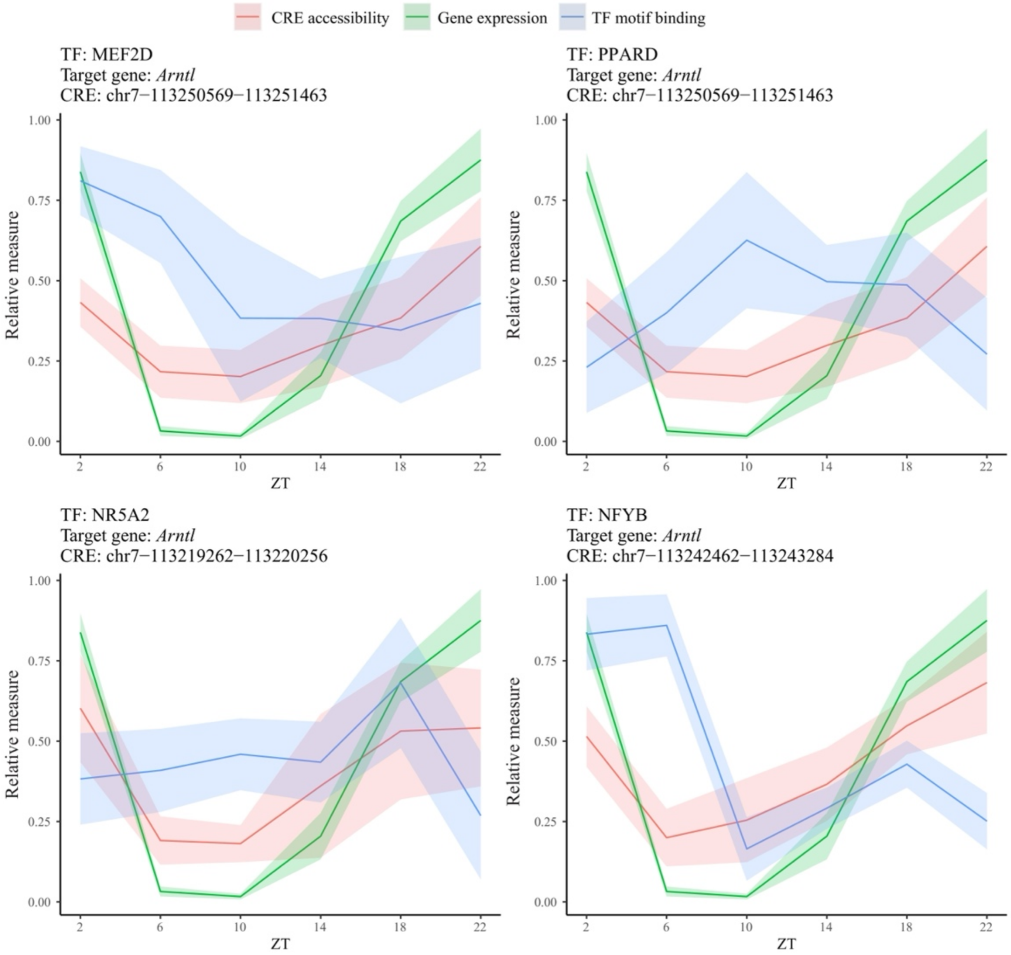

**Fig. S26.** Gene regulatory networks of core clock genes, categorized by E-box, D-box, and RORE motifs, as well as those without an annotated motif. Arrows indicate regulator-gene relationships, with colors denoting validation by orthogonal omics measurements.

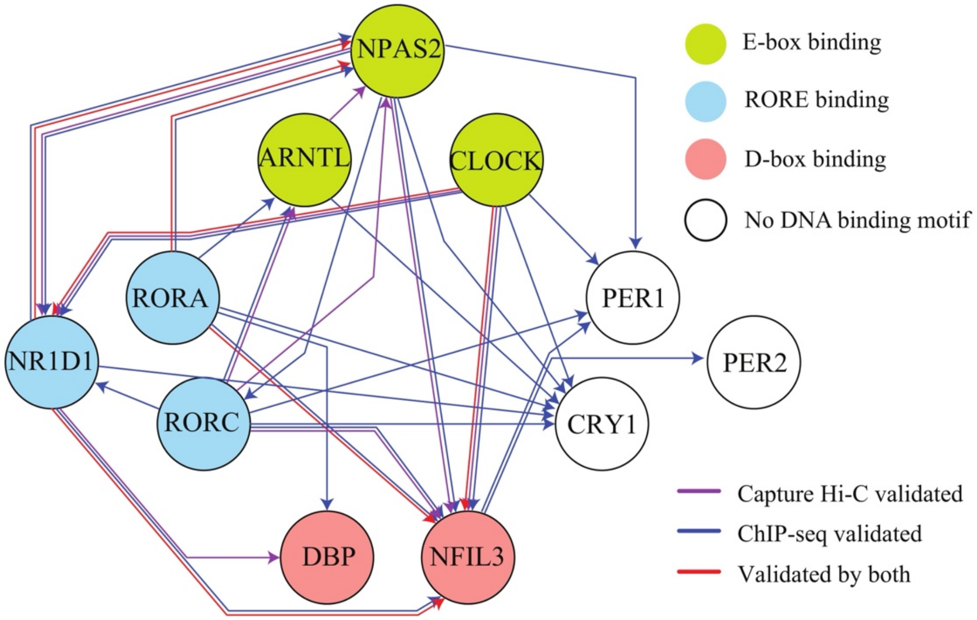

**Fig. S27.** TSS enrichment of ATAC fragments at ZT2 – ZT6, validated by time-series ChIP-seq data from previous studies [16,17]. (A) TSS enrichment from this study. (B) ChIP-seq of RNA polymerase II (Pol II). (C) ChIP-seq of TBP (TATA box binding protein), which is responsible for Pol II loading and initiation. (D) ChIP-seq of NELF-A, which is responsible for stabilizing Pol II in pause state. (E) H3K4me3 and (F) H3K36me3 serve as positive and negative histone modification markers for promoters near the TSS [17].

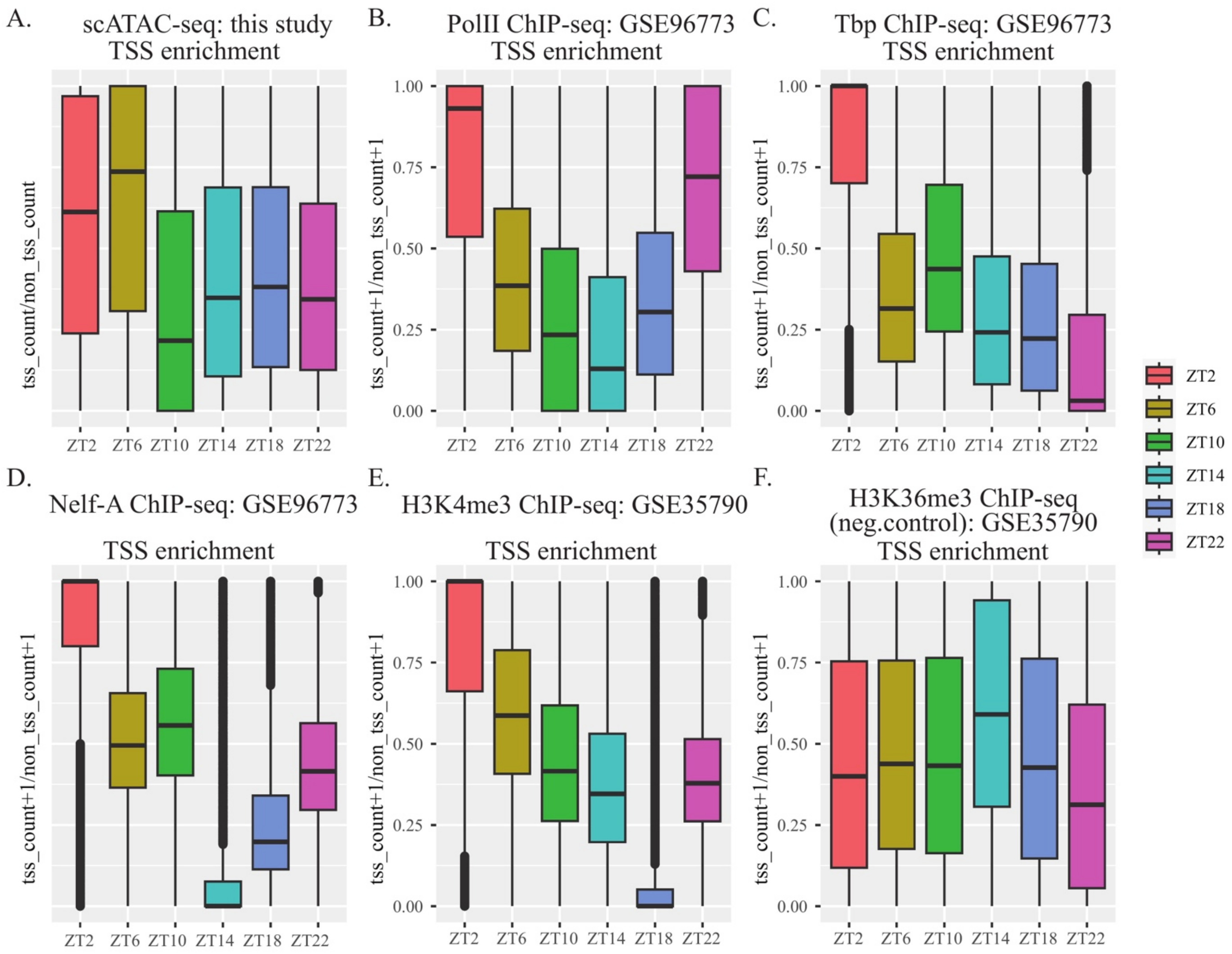

**Fig. S28.** RNA smFISH signals by Gaussian mixture model. (A) *Arntl*. (B) *Per1*. R package mclust is used with number of clusters set at two. Because exonic probes were used in the smFISH experiment, only larger dots that colonize with the genes from the cell nuclei were used in the calculation; we did so by keeping the cluster with the larger mean to enhance signal-to-noise ratio.

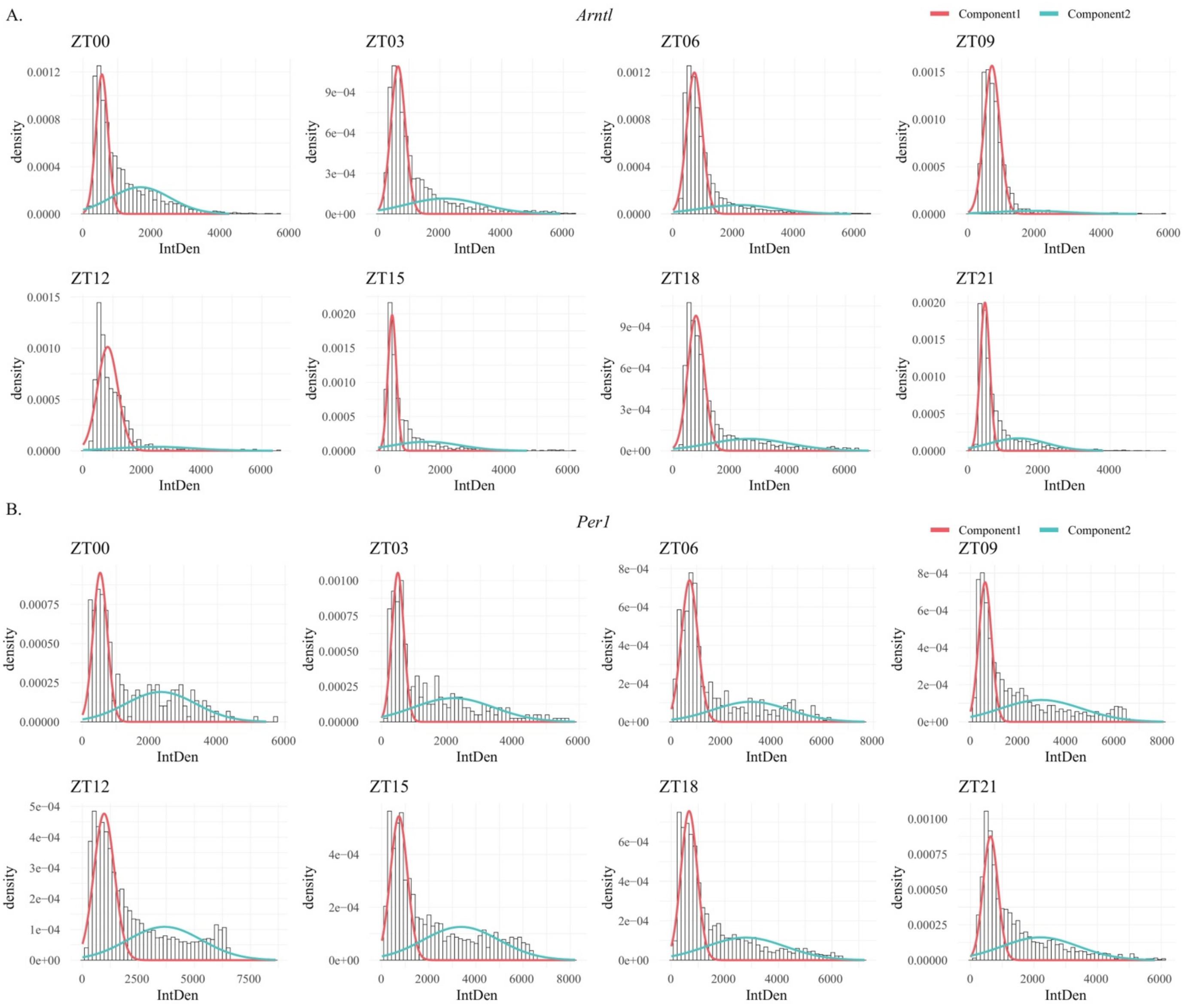

**Table S1.** Cell-type-specific circadian rhythm testing results of hepatocytes, endothelial cells, fibroblasts, and Kupffer cells. Significant rhythmicity is determined using the Cauchy combination test, with a Benjamini-Hochberg adjusted q-value (BH.Q) threshold of less than 0.01. Cells are downsampled to an equal count for each cell type to ensure the same testing power. Separately attached as an xlsx file.

**Table S2.** Rhythmic TF gene expression, ATAC activity, and motif scores in hepatocytes. Significant rhythmicity is determined using the Cauchy combination test, with a fold change threshold (TF gene expression fold change > 1.6, ATAC activity fold change > 1.1, motif scores fold change > 1.1) and a Benjamini-Hochberg adjusted q-value (BH.Q) threshold of less than 0.01. Separately attached as an xlsx file.

**Table S3.** Spatial, rhythmic, and spatial & rhythmic CREs identified via linkage analysis associating CRE accessibilities with target gene expressions across location and/or time in hepatocytes. Separately attached as an xlsx file.

**Table S4.** Meta output of hepatocyte's TRIPOD results which are validated by TF ChIP-seq or promoter capture HI-C, Significant trios is determined using Benjamini-Hochberg adjusted q-value (BH.Q) threshold of less than 0.01. Separately attached as an xlsx file.
